# Benchmarking the impact of reference genome selection on taxonomic profiling accuracy

**DOI:** 10.1101/2025.02.07.637076

**Authors:** Jasper van Bemmelen, Ioanna Nika, Jasmijn A. Baaijens

**Affiliations:** Intelligent Systems Department, Delft University of Technology, Van Mourik Broekmanweg 6, Delft, 2628 XE, Zuid-Holland, Netherlands; Department of Biomedical Informatics, Harvard Medical School, Shattuck St. 10, Boston, 02115, Massachusetts, United States

**Keywords:** Taxonomic profiling, Lineage abundance estimation, Sequence dereplication, Sequence clustering, Genome selection

## Abstract

**Background:** Over the past decades, genome databases have expanded exponentially, often incorporating highly similar genomes at the same taxonomic level. This redundancy can hinder taxonomic classification, leading to difficulties distinguishing between closely related sequences and increasing computational demands. While some novel taxonomic classification tools address this redundancy by selecting a subset of genomes as references, insights regarding the impact of different reference genome selection methods across taxonomic classification tools are lacking.

**Results:** We systematically evaluate genome selection and dereplication methods on bacterial and viral datasets using simulated metagenomic samples. For bacterial species-level profiling, incorporating all available genomes generally yields the highest accuracy, while having a limited impact on computational resource usage. In contrast, for highly similar bacterial strain-level and SARS-CoV-2 lineage-level datasets we find that selection significantly improves abundance estimation accuracy. Incorporating location-based metadata further enhances viral profiling performance by prioritizing locally relevant genomes. Across viral experiments, smaller reference sets significantly reduce memory and runtime requirements during both indexing and profiling, although this comes at an additional pre-processing cost.

**Conclusions:** Reference genome selection influences both accuracy and computational efficiency in taxonomic profiling, but its benefits seem context- and resolution-dependent. Our results demonstrate that reference set design does not have a one-size-fits-all solution, and that selection strategies should be adapted based on the biological and computational setting.

## Introduction

Taxonomic profiling forms one of the cornerstones of metagenomic analyses, allowing researchers to characterize and contextualize metagenomic samples by comparing samples against a database with a known taxonomy. Over the last decades, we have seen an explosion in the availability of genomic data, with databases hosted by the National Center for Biotechnology Information (NCBI) [55] and the Genome Taxonomy Database (GTDB) [43] doubling in size every few years. As taxonomic profiling inherently relies on the use of genome databases, this means that taxonomic profiling tools have to cope with their explosive growth.

In recent years, many new taxonomic profiling tools and pipelines have been developed. Several of these explicitly address the consequences of ever-increasing genome databases. For example, Kraken2 [56], one of the most popular taxonomic profiling tools, efficiently deals with large taxonomic databases by representing genomes as minimizers (a subset of the constituent *k*-mers of a sequence). This significantly reduces the storage and memory requirements compared to its predecessor. In a similar fashion, CLARK [42] only stores *k*-mers that are unique to the genomes in the database. Instead of using a compressed *k*-mer-based representation, Centrifuge [25] progressively builds a pangenome for every species by adding non-redundant segments of genomes. Marker-based taxonomic profiling tools, like MetaPhlan [7] and mOTUs [48], use databases which only store marker gene sequences. All of the profiling tools above focus on compressed representations of a database, rather than sequence selection.

Taxonomic profiling can also benefit from excluding redundant or less informative sequences through careful selection of representative genomes. Besides improving scalability, this approach can enhance profiling accuracy. Genome databases have taken a first step by selecting curated, high-quality genome sequences. A primary example is NCBI’s RefSeq [45], one of the most widely used genome databases, which has a selection of representative genomes for over 28,000 named species over all taxonomic kingdoms [19] and serves as a standard for many taxonomic profiling tools. Similarly, GTDB, which currently hosts over 600,000 microbial genomes, also provides a set of high-quality (i.e., low contamination, high completeness) reference genomes. Its representative genomes define species boundaries within the taxonomy and are updated as higher-quality genomes become available [43]. These curated databases play an important role in enabling accurate and efficient taxonomic profiling.

However, our understanding of optimal strategies for genome selection in the context of taxonomic profiling is limited and further selection may be desired. Recent methods [14, 27] incorporate a selection step to filter out highly similar genomes during index generation, which reduces false positive rates [27]. This step, often called sequence dereplication, selects a small subset of sequences such that every sequence is represented by at least one selected sequence. Tools and software suites that do so include CD-HIT [16], UCLUST [15], MeShClust [18] and MMseqs2 [51], and were primarily developed to dereplicate many short sequences such as reads. It remains unclear how widely applicable such selection strategies are and to what extent they can improve taxonomic profiling pipelines through reference genome selection.

While several benchmarking studies have compared the performance of different taxonomic profiling tools ([33, 36, 38, 49, 50]), these rely on a fixed selection of reference genomes to ensure fair comparisons across methods. Some studies have explored the effects of different reference databases, but only for different versions of RefSeq [39] or other standard databases in combination with different profiling parameters [57, 58]. Other work has focused on the use of taxon-specific reference sets, highlighting how this can bias results [46], or how dereplication tools scale in the context of short sequence dereplication (e.g., reads) [23, 54, 61]. However, to the best of our knowledge, no studies have systematically examined how variations in reference selection—beyond database versions or taxon-specific choices—affect the performance of taxonomic profiling tools.

To address this gap, we assess the impact of different genome selection techniques as a pre-processing step in taxonomic profiling. We consider a global database of potential reference genomes and apply a range of sequence dereplication methods, including VSEARCH [47], MeShClust [18], Gclust [29], and GGRaSP [11], as well as hierarchical clustering approaches. These methods aim to filter out sequences that are highly similar to other sequences, thereby framing the problem of reference genome selection as a specific instance of sequence dereplication. Using the filtered reference sets generated by these methods, we construct taxonomic profiling indexes and benchmark their effect on profiling accuracy in simulated metagenomic samples and a bacterial mock community. We find that, when target genomes are highly similar, using hierarchical clustering to select reference genomes can significantly improve taxonomic profiling accuracy while reducing computational resource requirements. When target genomes are less similar, we instead find that more sophisticated dereplication methods like MeShClust with a high similarity threshold can improve accuracy. This highlights the value of reference genome selection in optimizing taxonomic profiling.

### Overview of sequence dereplication methods

Sequence dereplication tools aim to reduce redundancy in a set of sequences by finding a subset of representative sequences. To do so, they address the following coverage-based problem: given a set of sequences *S*, find a subset *S^′^* ⊆ *S* that is representative of *S*. Although definitions of representative can depend on the dereplication method, the most commonly considered definition defines a subset *S^′^* ⊆ *S* to be representative of *S* if, for a given similarity function *f* and similarity threshold *T*, it holds that for every *s* ∈ *S*, there exists at least one sequence *s^′^* ∈ *S^′^* such that *f* (*s, s^′^*) ≥ *T*. The key idea in this formulation is that every sequence in *S* is represented by some sequence in *S^′^*. Below, we describe several categories of existing dereplication methods.

### Greedy incremental clustering

Greedy incremental clustering methods have been developed to operate on large numbers of sequences, making them applicable to modern day sequence volumes. Since comparing sequences based on exact alignment is computationally expensive [1], and the number of comparisons scales quadratically with the number of sequences, these tools focus on minimizing the number of sequence comparisons. In the greedy incremental clustering paradigm, sequences are sorted according to a total order (often from longest to shortest) and the first sequence is selected as a representative of its own cluster. Afterwards, subsequent sequences are compared to existing representatives, adding them to an existing cluster if their similarity with the corresponding representative is larger than the given threshold *T*. If no such sequence exists among the currently selected representatives, the sequence instead becomes a representative of its own cluster.

Many popular sequence dereplication tools, like MMseqs2 [51], CD-HIT [16], UCLUST [15], VSEARCH [47] and Gclust [29] utilize the greedy incremental clustering framework. These methods have distinct implementations which differ primarily in how they define the similarity function *f*, or in how they apply filters to reduce the number of sequence comparisons. For example, CD-HIT performs global alignment and then defines the similarity as the number of identical characters divided by the length of the shortest sequence of the two, whereas UCLUST divides by the average length of both sequences. More recently, the LINCLUST algorithm was added to the MMseqs2 software suite. LINCLUST finds groups of sequences sharing at least one *k*-mer, defining the longest sequence of each group as its center, and merges groups that share a center so that alignment-based comparisons only have to be done between sequences and the centers of groups they belong to [52]. This approach was shown to scale approximately linear with the number of input sequences, making it feasible to scale to billions of sequences.

### Submodular optimization

In addition to the greedy incremental clustering paradigm, there are also methods that consider a coverage-based problem formulation, but instead solve it differently. In Repset [31], it is shown that problems adhering to a coverage-based selection formulation are related to unconstrained problems in which a submodular function is optimized instead. By using known approximation algorithms for optimizing submodular functions, [31] shows that this dereplication works well for dereplicating protein sequences. Furthermore, they also show that extensions can be incorporated with allows for bi-objective functions in which the number of selected representatives is explicitly minimized. However, the authors also show that, compared to greedy incremental clustering methods, their approach is significantly slower, making it less applicable to large scale datasets with millions of sequences.

### Mean-shift clustering

Mean-shift clustering provides an alternative approach where cluster representatives are iteratively shifted towards the center of mass of their respective clusters. MeShClust [18] is based on a mean-shift [13] algorithm where pairs of clusters are merged when the pairwise similarity of their representatives exceeds a similarity threshold. Representatives are re-selected every iteration as the sequence closest to the center of their respective clusters. Therefore, this approach finds more meaningful clusters and cluster representatives compared to approaches that fix representatives, by allowing cluster representatives to be updated [22]. Additionally, MeShClust incorporates a generalized linear model that is trained to classify pairs of sequences as being similar according to the provided threshold *T*. While this requires training a model upfront, it results in relatively fast sequence comparisons during the mean-shift algorithm. Hence, MeShClust can run on thousands of full-length bacterial genomes (∼3Mbp) despite the computational complexity of the mean-shift algorithm [18].

### Hierarchical clustering

Another popular strategy for sequence dereplication is to apply hierarchical clustering, after which representatives per cluster are selected. This approach is implemented by both dRep [40] and GGRaSP [11], and is also used as a pre-processing step by recent taxonomic profilers such as StrainGE [14] and YACHT [27]. GGRaSP clusters sequences using either a user-defined similarity threshold *T*, or a threshold inferred from the data using Gaussian mixture models. It selects the medoid sequence of every cluster as a representative, but also allows for a ranking of the input sequences to prioritize the selection of sequences that meet certain user requirements (e.g. assembly completeness). dRep follows a similar strategy but employs a two-step procedure: first a rough clustering of the input sequences, then hierarchical clustering within every preliminary cluster using the user-defined similarity threshold *T*.

Hierarchical clustering-based methods require all-vs-all similarity calculations, which are computationally expensive. Fast similarity estimators like MASH [41] and fastANI [21] are therefore essential in practice. In dRep, the first clustering step relies on MASH distances, while the second step uses fastANI; the latter is more accurate but slower than MASH, thus applying fastANI only within preliminary clusters reduces the number of expensive comparisons.

### Alternative dereplication methods

Several methods have been developed to perform sequence dereplication under more specific modeling assumptions. PARNAS [35], for example, treats the dereplication problem as a *k*-medoids problem or optimal coverage problem on a phylogenetic tree. By requiring a phylogenetic tree structure, distances between sequences adhere to a tree-metric, which makes the corresponding dereplication problems computationally feasible. Another approach, TARDiS [34], uses a genetic algorithm to select a pre-defined number of sequences while optimizing for maximal sequence diversity and temporal distribution. This enables incorporation of sampling time as metadata when available, but requires the user to specify the desired number of representative sequences beforehand.

In contrast to explicitly selecting representative sequences through dereplication, ReprDB and panDB [60] were introduced as a method to find a compressed representation of the pangenome of a species. These methods work by starting with an initial representative genome, followed by iteratively aligning genomes from the same taxonomic group (i.e. target) in order to detect novel genomic fragments that are added to the pangenome representation, similar to the approach used by Centrifuge. Although such methods allow for a compressed representation of a set of genomes, they do not select representatives genomes directly.

## Methods

The general reference set construction workflow involves three steps: genome retrieval, optional quality filtering or downsampling, and sequence dereplication (Figure 1). This section provides an overview of the tools used, followed by a description of datasets, benchmarking set-up, and performance evaluation.

**Fig. 1.**
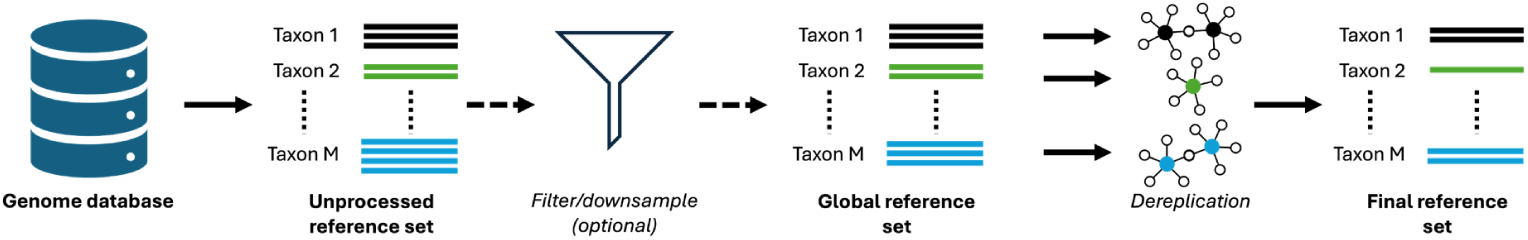
A schematic representation of the reference genome selection pipeline. Potential reference genomes are first retrieved from a database. Afterwards, quality filtering and/or downsampling can be applied. Finally, sequence dereplication methods are applied to the genomes of every taxon of interest (individually), resulting in a final reference set.

### Selection of dereplication tools

For our experiments, we selected dereplication tools representing the main methodological categories described above.

From the greedy incremental clustering family, we included VSEARCH v2.28.1 and Gclust v1.0. UCLUST was omitted since its unrestricted 64-bit version is only available as commercial software while having similar functionalities as VSEARCH. Gclust, which uses sparse suffix arrays in order to accelerate sequence similarity searches in the incremental clustering framework, was included as an efficient alternative to VSEARCH. Tools from the MMseqs2 suite were excluded as they failed to produce results while setting up experiments (see Supplementary Material). Additionally, CDHIT and the submodular optimization-based method Repset were excluded due to prohibitive runtime requirements in preliminary experiments. As these omitted methods operate on a similar basis, we expect that they would have produced reference sets highly similar to those generated by Gclust and VSEARCH.

As a representative of mean-shift clustering approaches, we included MeShClust, using the latest available version (Identity v1.2, MeShClust v2.0), which has been shown to produce clusters of higher quality than its predecessor [18].

For hierarchical clustering-based approaches, we estimated genome distances using MASH v2.3, and included GGRaSP v1.3, which automatically determines similarity thresholds using Gaussian mixture models, and a custom implementation based on the SciPy library v1.5.0 (https://scipy.org). The latter allows reuse of the precomputed similarity matrix generated for GGRaSP, whereas dRep computes similarities at runtime. Since our reference genome pool was already filtered based on quality metrics similar to those used by dRep, its additional scoring procedure was not required.

For the viral experiments, we also evaluated the Viral Lineage Quantification (VLQ) selection pipeline [3], a selection strategy developed for SARS-CoV-2 that greedily selects genomes containing mutations occurring at a frequency of at least 50% relative to a standard reference genome.

Finally, PARNAS was excluded since it requires the input to adhere to a treemetric, and TARDiS was not considered as our experiments do not explicitly incorporate temporal metadata.

### Datasets

We designed our experiments to consider two main scenarios: bacterial genomes and viral genomes (specifically SARS-CoV-2). For the bacterial scenario, the focus is on taxonomic classification at the species- and strain-level, whereas for the viral scenario, the goal is to estimate abundances at the lineage level. This set-up allows us to investigate how the impact of reference genome selection differs between datasets with varying degrees of sequence similarity. We focus on impact in terms of taxonomic profiling accuracy, but we also report the efficiency of the dereplication, index construction, and taxonomic profiling in relation to the different dereplication tools used.

#### Bacterial experiments

For our bacterial analysis, we distinguish between species-level and strain-level profiling based on NCBI’s taxonomy (downloaded September 24th, 2024). For the species-level experiments we considered mixtures of five *Streptococcus* species: *Streptococcus suis*, *Streptococcus thermophilus*, *Streptococcus agalactiae*, *Streptococcus pneumoniae* and *Streptococcus pyogenes*. These species were selected to maximize diversity within the Streptococcus genus (see Supplementary Material). Strain-level experiments were performed using *Escherichia coli* strains to evaluate performance at increased resolution. The included strains were: H10407, UTI89, Sakai and E24377A, based on the mock community design (see below).

For species-level experiments, we performed dereplication at the species-level, and for the strain experiments at the strain-level (Figure 1), using MeShClust, Gclust, GGRaSP and a custom implementation of hierarchical clustering-based selection. The reference genomes used throughout all experiments consisted of all “Complete Genome” status assemblies from NCBI Assembly (now NCBI Genomes), downloaded on October 9th, 2024. We only included species or strains that had at least one such assembly. After dereplication, we performed taxonomic profiling at the corresponding level, to determine the resulting profiling accuracy for each dereplication strategy at different resolutions.

Finally, to further assess how the impact of reference genome selection varies with target similarity, we also performed species-level experiments for mixtures of species from the same family and order, respectively (see Table S1 in Supplementary Material). Results for the order- and family-based species-level experiments were highly consistent with those in the genus-based species-level experiments, and all details and results are provided in the Supplementary Material.

##### Dereplication parameter settings

In the species-level experiments, all tools were run with similarity thresholds of 95%, 97% and 99%, except for GGRaSP for which we used the Gaussian mixture model based threshold selection strategy. The 95% similarity threshold reflects the cut-off for species definitions in prokaryotes [26], and the higher thresholds (97%, 99%) may create more refined reference sets. For the strain-level experiments, we ran tools with similarity thresholds of 95%, 99% and 99.9% as strains are more closely related. For MeShClust, the maximum allowed similarity threshold is 99%, so we used only the 95% and 99% thresholds. If a method failed to produce a selection for a species or strain, we selected a random reference genome to represent it.

For both GGRaSP and the other hierarchical clustering-based selections we ran MASH to estimate distances between all pairs of genomes within a species. Here we used the default *k*-mer size of 21 and sketch size of 1000 for the species-level experiments, and *k* = 31 with a sketch size of 5000 for the strain-level experiments. For the hierarchical clustering selections we generated selections using both single-linkage clustering and complete-linkage clustering at the designated thresholds. Representatives for each cluster were obtained by selecting the medoid of respective clusters.

To compare against a naive selection strategy, we also generated a medoid reference set containing only the medoid genome of every species or strain (based on MASH distances). Moreover, as a baseline, we include the “All” reference set, which contains all genomes (no selection). The exact commands (including parameter settings) to produce all reference sets can be found on Github [5], and the number of threads used for every method are given in Supplementary Tables S2–3.

##### Index construction and taxonomic profiling

For profiling we use Bracken v1.0.0 [32] (building on Kraken2 v2.1.3 [56]), Centrifuge v1.0.4.2 [25] and DUDes v0.10.0 [44] (using BWA v0.7.18 [28] for alignment). This selection of methods covers two popular *k*-mer based profiling tools with different approaches for compressing reference genomes (Bracken, Centrifuge) as well as one exact alignment based tool (DUDes). For all methods, reference indexes were constructed using default parameters, except for the number of threads (Supplementary Tables S2–3).

One complicating factor is that these taxonomic profiling tools do not estimate taxon abundances in the same way. Bracken estimates “sequence abundance”, which is defined as the number of reads assigned to a particular taxon divided by the total number of reads that have been classified. In contrast, DUDes estimates “taxonomic abundance”, which is defined as the relative number of occurrences of a particular taxon in a sample. The core difference between these abundance definitions is that taxonomic abundances account for genome lengths and thus these outcomes cannot be compared directly [53]. Centrifuge is able to estimate both sequence abundance and taxonomic abundance, and we evaluate accuracy based on estimated sequence abundances derived from uniquely mapped reads.

##### Read simulations

To assess the performance of taxonomic profilers using different reference sets, we simulated paired-end Illumina reads for every experiment using ART v2016.06.05 [20]. For all bacterial experiments, we generated reads with a length of 150 basepairs (bp), and a mean fragment size of 270bp with a standard deviation of 20bp. Furthermore, for every experiment we simulated 10 metagenomic samples with approximately 4,500,000 HiSeq2500 readpairs, generated from genome assemblies from NCBI Assembly (see Supplementary Material for details).

##### Strain mock community samples

For the strain experiments we included an additional set of experiments where we profile the *in vitro* mock community sample (SAMN17091845) described in [14]. This sample consists of 99% human host DNA and 1% *E. coli* DNA distributed over the four *E. coli* strains as follows: 80% H10407, 15% UTI89, 4.9% Sakai and 0.1% E24377A. Before profiling, reads were checked with FastQC v0.12.1 [2], after which adapters were removed and low quality bases (quality threshold below 20) were trimmed from read tails using FastP v1.1.0 [10]. Reads shorter than 50bp were discarded after trimming, and remaining reads were mapped to the GRCh38 human reference genome using Bowtie2 (v2.5.4) with the –very-sensitive preset. Readpairs that did not align concordantly were extracted using samtools v1.23 (-f 12 -F 256), resulting in a collection of 726,777 readpairs.

In addition to the real sample, we also simulated 10 analogous artificial HiSeq2500 samples using ART with a mean fragment size of 265 and standard deviation of 104 (estimated from the real sample using Picard v2.20.4 [9]), which replicate the mock community experiment while providing a hard ground truth.

#### Viral experiments

For the viral experiments, we focused on SARS-CoV-2 to evaluate the effect of reference genome selection on lineage-level abundance estimation. We created simulated samples representing SARS-CoV-2 populations detected in wastewater at a specific point in time in Connecticut, USA. Similar to the bacterial experiments, we performed 3 sets of experiments, differing in the geographic filtering applied to the genome database (GISAID [24]): i) global experiments, with no geographic filter; ii) country experiments, including only reference genomes from the same country (USA) as the samples; and iii) state experiments, including only genomes from the same state (Connecticut) as the samples. This set-up allows us to explore whether incorporating geographic information into reference genome selection can improve lineage-level abundance estimation, an aspect particularly relevant for environmental samples such as wastewater. In each experiment, we performed sequence dereplication at the lineage level (Figure 1). We used the VLQ pipeline [3] to perform lineage-level abundance estimation and evaluate abundance estimation accuracy similar to the bacterial experiments.

##### Dataset composition

The genomes used throughout the viral experiments were downloaded from GISAID [24] on June 12, 2022, and consisted of sequences from the period January 1st 2021 until March 31st 2021, satisfying a genome length of at least 25,000bp, and an N-content of at most 0.1%. After geography-based selection, genomes were randomly downsampled to at most 1000 sequences per lineage to keep computations feasible.

##### Parameter settings

For both GGRaSP and the other hierarchical clustering methods, we used MASH to estimate pairwise sequence similarities, using a sketch size of 5,000 and *k*-mer size of 31 (similar to the bacterial strain-level experiments). We ran GGRaSP with the automatic threshold selection strategy. For the single-linkage and hierarchical-linkage clustering selections we calculated the 1st, 5th, 10th, 25th, 50th, 90th and 99th percentile distances (based on MASH distances) for every lineage individually, and used these as similarity thresholds. VSEARCH, Gclust and MeShClust define similarities during selection, so for these tools we used pre-defined similarity thresholds of 95%, 99%, and 99.9% (SARS-CoV-2 genomes are generally more than 99% similar), again excluding the 99.9% similarity threshold for MeShClust. For the selection with VLQ we used a minimal allele frequency threshold of 50%, which ensures that all point-wise mutations (compared to the MN908947.3 reference genome) that occur with a frequency of at least 50% (within their respective lineage) are captured in the selected genomes.

The exact commands used to run every tool can be found on Github [5], and the number of threads used for every method are given in Supplementary Tables S4–6. In the event that a method failed to select one or more representatives for a given lineage, we randomly select a representative from that lineage. Finally, we consider a medoid and “All” reference sets, similar to the bacterial setting.

##### Index construction and taxonomic profiling

For viral profiling we use the VLQ pipeline [3] (which uses kallisto [8], v0.44.0). Both index construction and subsequent abundance estimation were done using default parameters, and abundance estimations were further processed using the scripts available in the VLQ pipeline that convert kallisto output (transcripts per million) to relative abundances at the lineage level.

##### Read simulations

For the viral experiments we simulated paired-end Illumina HiSeq2500 reads now using SARS-CoV-2 genomes having Connecticut as their marked location, and April 30th, 2021 as their sampling date. This constitutes a collection of 25 genomes used for simulations, spread across 6 lineages. We simulated 20 samples of approximately 9,000-10,000 paired-end reads with a length of 150bp, and a mean fragment size and standard deviation of 250bp and 10bp respectively. Since preliminary experiments showed large effects induced by reference genome selection, we repeated these experiments fixing the relative abundance of lineage B.1.1.7 (commonly known as the alpha variant) at 1%, 10%, 20%, 30%, 40%, 50%, 60%, 70%, 80%, 90% and 100%. Background sequences were randomly simulated from other lineages, and this set-up allows us to assess whether the effect of reference genome selection on profiling accuracy holds at different sample compositions.

### Performance evaluation

All experiments presented in this work were run on the Delft Artificial Intelligence Cluster (DAIC) high performance computing cluster [12]. The benchmarking pipeline and commands are publicly available on Github. In what follows we will describe how we compare experimental outcomes.

#### Reference set comparisons

In all experiments, we calculate containment indices (CI) between every pair of reference sets. Formally, the CI is defined as 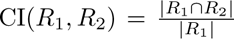, where *R*_1_ and *R*_2_ are sets of reference genomes. For our results, we only consider genomes from taxa that have at least 2 sequences to calculate containment indices, since otherwise the intersection for a given taxon is always equal to 1. Similar to the Jaccard index, the containment index quantifies the similarity of sets. Unlike the Jaccard index, however, the containment index is asymmetric in its arguments, providing additional information when reference sets differ in size.

We assess statistical significance of the observed containment indices using Monte Carlo simulations, as described in the Supplementary Material. Since multiple containment indices were tested (15×15 for the bacterial species-level experiments, 14×14 for the bacterial strain-level experiments, and 26 × 26 for viral experiments), we corrected for multiple hypothesis testing using the Benjamini-Hochberg procedure applied to all off-diagonal p-values excluding comparisons versus the “All” reference set. Adjusted p-values below 0.05 were considered statistically significant.

#### Taxonomic profiling evaluation

To evaluate the accuracy of taxonomic profiling we calculate both the L1-norm and F1-score of abundance estimates compared to the ground truth for every profiler, sample and reference set. As in OPAL [37], we define the L1-norm of predicted abundances 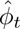 for all taxa *t* as:

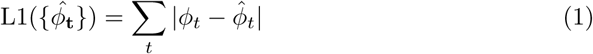

where *ϕ_t_* denotes the ground truth abundance of taxon *t*. The L1-norm measures the total error made between the true and predicted abundances, and ranges from 0 to 2. For simplicity, we transform the computed L1-norms in the following way:

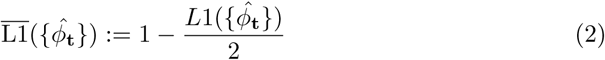

to obtain what we will denote the *abundance accuracy*. The abundance accuracy is equal to one minus the Bray-Curtis dissimilarity for abundance estimates and ranges from 0 to 1, where a value of 1 indicates a perfect estimation of all abundances, and a value of 0 indicates a totally incorrect estimation of abundances.

The F1-score is defined as the harmonic mean of the precision and recall of estimations, and ranges between 0 and 1. A value of 1 indicates that both the precision and recall are perfect, meaning that every taxon with nonzero abundance in the ground truth has a nonzero estimated abundance, and every taxon with zero ground truth abundance also has zero estimated abundance.

We reduce false positives by removing any taxa with predicted relative abundance below 0.1% (0.05% for bacterial strain-level experiments), and subsequently re-scale remaining abundances to sum up to 100%. For all experiments, profiling methods, and reference sets, we compare both the abundance accuracy and F1-score to the respective scores of the “All” selection. We test whether a selection achieves significantly higher accuracy than the “All” selection (for both abundance accuracy and F1) by performing a Wilcoxon signed rank test, adjusting for false discovery rate using Benjamini-Hochberg correction within every experiment and for every profiler and metric separately. Adjusted p-values below 0.05 were considered statistically significant.

## Results

### Reference set compositions overlap across dereplication methods

To gain insight into reference set similarity across different dereplication tools, we calculated containment indices (see Methods) between reference sets. In calculating containment indices we ignored taxa (i.e. species or lineages) for which the database contained a single genome, as these are included in all reference sets by default.

Figure 2(A) shows that in the bacterial experiments, different reference sets have substantial overlaps with containment indices often exceeding 0.50. This effect is particularly prominent when comparing smaller reference sets (e.g. those selected through hierarchical clustering) to large reference sets, or when comparing different reference sets obtained from the same method using different similarity thresholds. Interestingly, the overlaps between all MeShClust and Gclust selections are high in general, despite their respective selection methodologies being largely distinct.

**Fig. 2.**
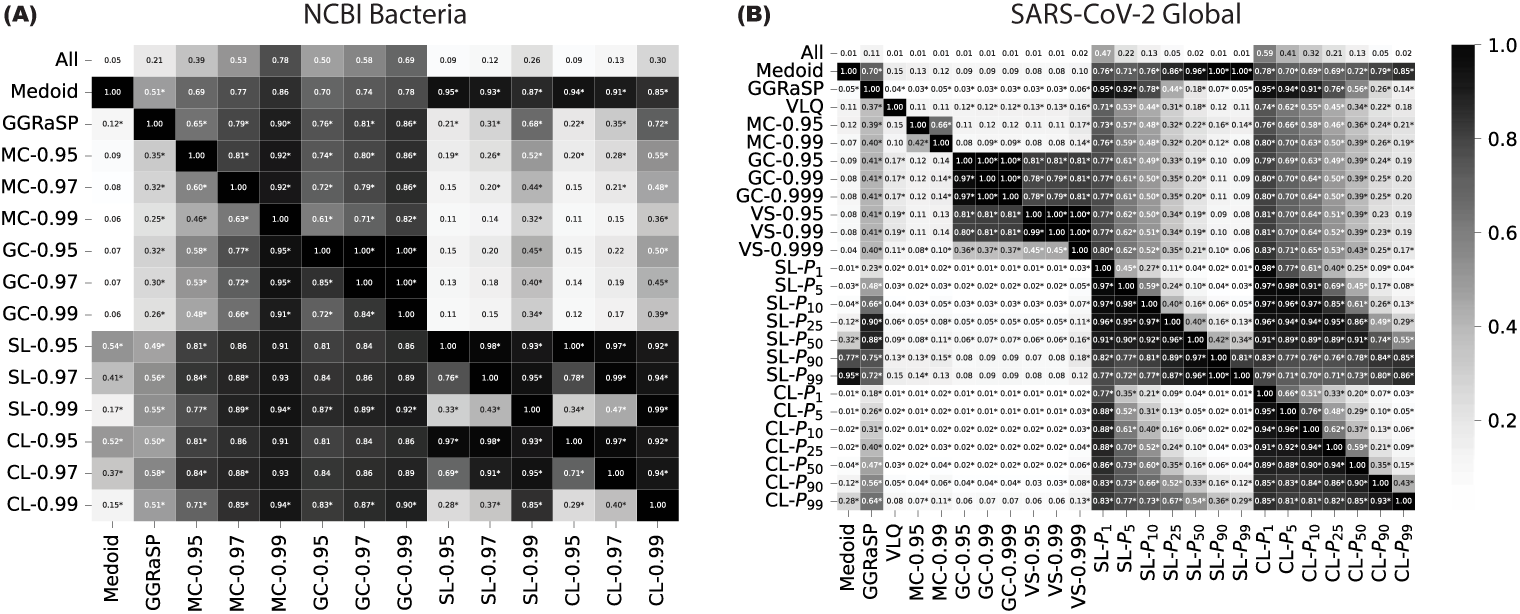
**(A)** Containment indices (CIs) between reference genomes selected by different dereplication tools for all bacterial genomes in NCBI, and **(B)** SARS-CoV-2 genomes (no filtering). CI is only calculated for taxa (species/lineages) that have more than 1 genome to select from. Asterisks indicate that the CI is significantly higher compared to randomly selecting genomes (*p <* 0.05, Benjamini-Hochberg corrected).

When comparing reference sets *R*_1_*, R*_2_ with |*R*_1_| ≫ |*R*_2_|, the containment index is generally small: the numerator in the containment index is bounded by the smaller reference set, while the denominator is determined by the larger reference set. Thus, to assess whether the containment indices are larger than expected by chance, we compare to an empirical null distribution and test for statistical significance (see Methods). In Figure 2, containment indices which are significantly larger than expected are indicated by an asterisk. We observe that even for small containment indices, their similarity is generally significantly larger than expected by chance. We observe the same overall trends for the reference sets limited to the taxa considered for the species-level experiments (Figures S1–3 in Supplementary Material). For the strain-level reference sets, we instead see that, generally, only containment indices between selections made by the same method are statistically significant (Figure S4 in Supplementary Material).

Viral dereplication resulted in small reference sets, particularly for methods using fixed similarity thresholds. The largest non-hierarchical clustering selection (VSEARCH-0.999) included only 2% of available genomes for lineages with multiple genomes available. In contrast, hierarchical clustering with dynamically determined thresholds yielded more variable reference set sizes, ranging from 1–2% to nearly 60% of all genomes. Because these reference sets are small, overlap between selections is limited (Figure 2(B)). Even so, containment indices were consistently higher than expected under random selection, indicating that distinct dereplication strategies do capture partially overlapping subsets of reference genomes.

To further quantify overlap, we assessed whether a “core selection” of reference genomes existed across methods (excluding the medoid selection). We measured this in two ways: (i) the number of genomes shared by all selections, and (ii) the number of species/strains/lineages containing at least one genome selected by every method (restricting analysis to selections that succesfully terminated for taxa). Despite statistically significant containment indices, no clear viral core selection emerged; only five genomes from distinct non-singleton lineages were shared by all methods (Figure S6 in Supplementary Material). In contrast, bacterial reference sets had 30-57% of non-singleton species or strains sharing at least one genome across methods (Figures S8–S12 in Supplementary Material). These results indicate that bacterial genome selections are relatively stable across dereplication methods, whereas viral selections are more sensitive to the underlying selection strategy.

### Reference selection increases profiling accuracy when targets are similar

To investigate the impact of reference genome selection on taxonomic profiling accuracy, we used the constructed reference sets to build taxonomic profiling indexes and analyze simulated metagenomic samples. We evaluate profiling accuracy by calculating abundance accuracy and F1-scores of every sample (see Methods).

#### Bacterial experiments

For each bacterial experiment, we obtained a sub-database by filtering out taxa that were not part of the selected taxon (see Supplementary Material). For the species-level experiments, we thus only retained genomes from all species of the *Streptococcus* genus, resulting in a sub-database of 1,502 genomes over 103 species from which we obtained 15 reference sets (one per method-threshold combination) that are used for downstream analyses. For the strain-level experiments, we only retain *E. coli* genomes with a strain annotation, yielding a sub-database of 203 genomes over 117 strains and 14 reference sets (MeShClust does not allow a threshold higher than 99% similarity). Figure 3 shows the abundance accuracy and F1-scores for the bacterial experiments, across taxonomic profilers. Across all experiments and profilers, accuracy scores have low variance, indicating consistent outcomes across samples. Furthermore, in the species-level experiments, the “All” reference set consistently ranks among the best performing selections, with median abundance accuracies between 0.8 and 0.9, and high F1-scores for all profilers.

**Fig. 3.**
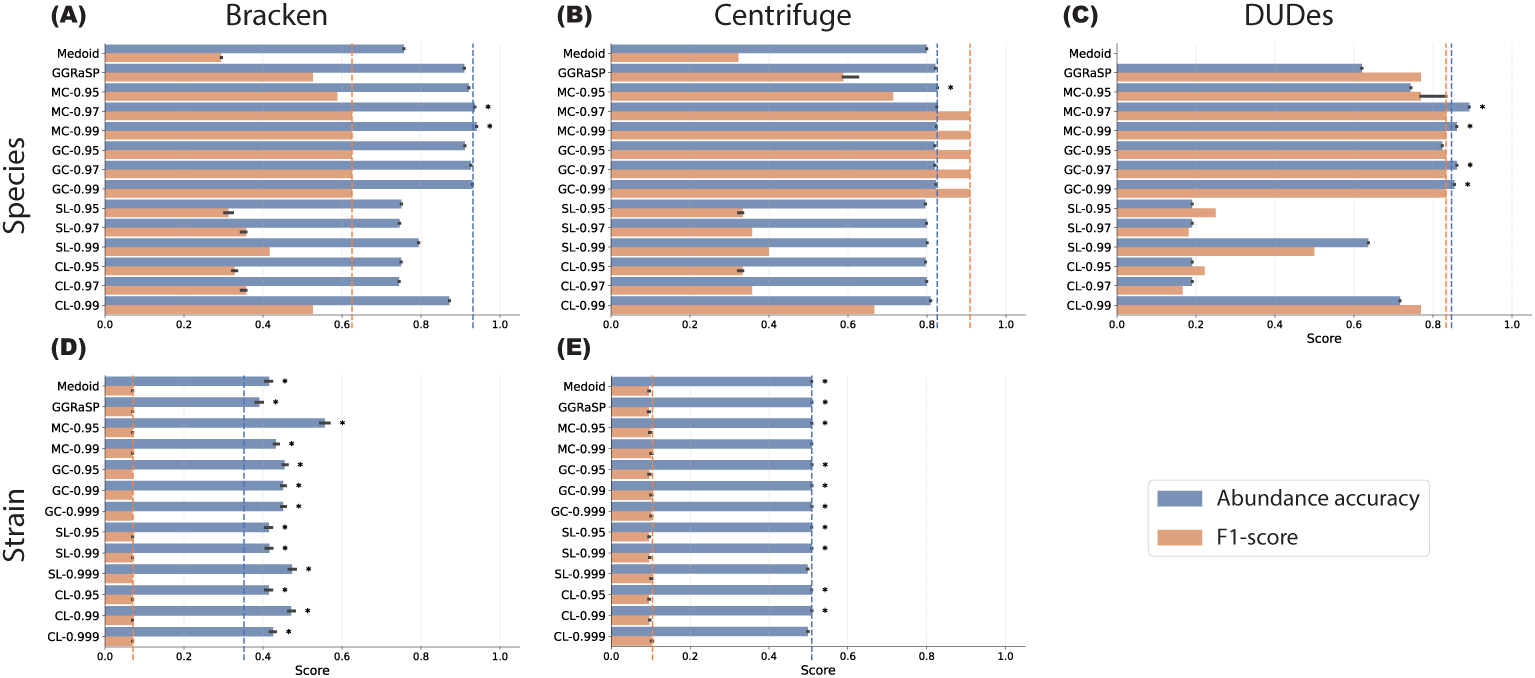
Accuracy metrics (abundance accuracy, F1-score) of taxonomic profiling tools in the bacterial experiments. Panels A-E are arranged in two rows (experimental setting: species-level, strain-level) and three columns (profiling tools: Bracken, Centrifuge, DUDes). Dashed vertical lines correspond to the median performance of the baseline “All” reference set (no selection) and error bars correspond to 10-90th percentile values. An asterisk indicates that the accuracy metric of a method is significantly better than that of the “All” reference set (Wilcoxon signed rank test, *p <* 0.05, Benjamini-Hochberg corrected). The panel for DUDes in the strain experiments is omitted as DUDes was unable to produce strain-level predictions. evaluate dereplication strategies when genetic diversity among reference genomes is extremely low.

In the strain-level experiments, abundance accuracy and F1-scores are generally much lower due to the increased similarity of target genomes, with DUDes even unable to assign abundance below species level for any reference set. However, in contrast to the species-level experiments, all dereplication methods result in better abundance accuracy scores compared to the “All” reference set when using Bracken. For Centrifuge, abundance accuracies are more similar across methods, with some methods marginally, yet statistically significantly, outperforming the “All” reference set. For both profilers, F1-scores are similar for all reference sets in the strain-level experiments, showing that in this setting the merit of reference genome selection lies in improving abundance estimation accuracy. We repeated both species-level and strain-level experiments in a low-coverage setting (450k reads instead of 4.5M reads), which showed increased variability in profiling accuracy, while exhibiting similar profiling accuracy relative to the “All” reference set (Figures S13–16 and Tables S7–10 in Supplementary Material).

No single combination of method and similarity threshold consistently outper-formed all others, and abundance accuracy does not seem to monotonously increase with similarity thresholds. Among dereplication approaches, MeShClust- and Gclust- based reference sets consistently rank among the best, often achieving comparable or better accuracy to the “All” reference set. These findings also hold for the family- and order-based species-level experiments, with outcomes relative to the “All” reference set being largely similar to those in the genus-based species-level experiments (Figures S13–16 in Supplementary Material). This suggests that carefully tuned reference genome selection significantly improves profiling accuracy when target genomes are highly similar, while maintaining similar profiling accuracy when target genomes are more divergent.

#### Viral experiments

To assess the impact of reference genome selection in a viral context, we focused on SARS-CoV-2 lineage-level abundance estimation. We used a database of 115,850 SARS-CoV-2 genomes spanning 1,020 viral lineages (see Methods), and we obtained 26 reference sets (one for each method-threshold combination) for downstream analysis. Since these genomes are generally more than 99 % similar, this set-up allows us to Figure 4(A) shows that several reference sets achieved significantly higher accuracy than the “All” reference set. Hierarchical clustering-derived reference sets (excluding GGRaSP) consistently performed best; the complete-linkage reference set with the 99th percentile similarity threshold achieved the highest median abundance accuracy (0.72 vs. 0.51 for the “All” set). Other clustering-based selections, including medoid, single-linkage, and complete-linkage sets at various thresholds, also significantly outperformed the “All” reference set in terms of abundance accuracy. By contrast, intermediate thresholds (25th and 50th percentile) for complete-linkage selections yielded no significant improvement. While F1-scores were generally low (a consequence of many false positives due to the high similarity of SARS-CoV-2 genomes), nearly all selections outperform the “All” reference set (median F1-score of 0.095). Overall, these results show that careful reference genome selection can significantly improve viral lineage abundance estimation accuracy.

**Fig. 4.**
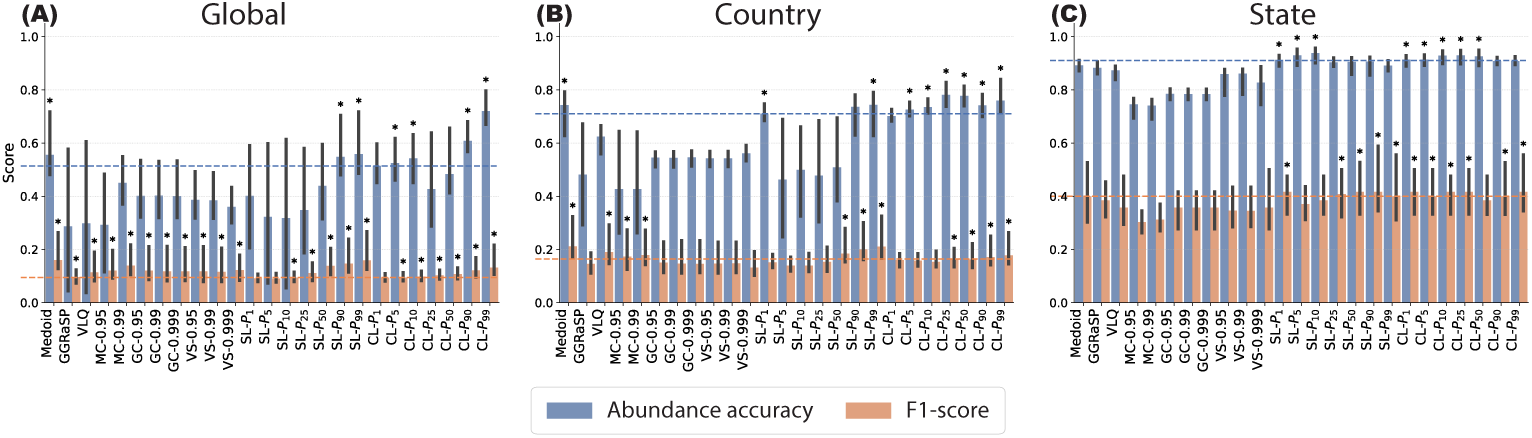
Accuracy metrics (abundance accuracy, F1-score) in the SARS-CoV-2 experiments. Panels A–C correspond to the three geographic filtering levels: **(A)** no filter, **(B)** country filter (USA), and **(C)** state filter (Connecticut). Dashed vertical lines correspond to the median performance of the “All” reference set (no selection) and error bars correspond to 10-90th percentile values. An asterisk indicates that the accuracy metric of a method is significantly better than that of the “All” reference set (Wilcoxon signed rank test, *p <* 0.05, Benjamini-Hochberg corrected).

The results for the viral experiments are consistent with the results for the bacterial strain-level results. One difference with the bacterial experiments, is that accuracies in the viral experiments are more variable, due to varying sample compositions. Additionally, methods that did well in the bacterial experiments (i.e. MeShClust and Gclust) perform substantially worse here, with conservative hierarchical clustering based selections generally performing best. This suggests that optimal reference set construction is context-dependent, and is primarily driven by the required resolution, as well as diversity and redundancy of underlying genomes. Moreover, these results show that selection strategies effective for bacterial profiling should not be directly transferred to viral datasets, and vice versa, without prior evaluation.

### Location-based reference selection improves profiling accuracy for SARS-CoV-2

Given the available metadata for the viral sequences, we investigated whether selecting references based on geographical location can further improve taxonomic profiling accuracy. Our viral samples were simulated to represent SARS-CoV-2 populations in wastewater at a specific point in time in Connecticut, USA. Thus, to evaluate the impact of location-based reference selection, we created reference sets at country and state level, including only sequences from the same country (USA) and state (Connecticut), respectively (see Methods). Following the same workflow as before, we estimated relative abundances of SARS-CoV-2 lineages to assess how geographic filtering affects profiling accuracy.

Figure 4**(B)** and **(C)** show the abundance accuracy and F1-score for all reference sets across all geographic filtering levels. Overall, both abundance accuracy and F1-score benefit from applying increasingly stringent filters: reference sets in the state-filtered experiments produce the most accurate results with the least variance. Notably, the average median abundance accuracy across all methods increased from 0.442 to 0.875 (+109%), with median F1-scores improved from 0.116 to 0.382 (+240%) on average when comparing state results versus baseline results.

The hierarchical clustering based selections remain consistently more accurate than the “All” selection after applying location filters. In the state-filtered experiments, selections with stricter similarity thresholds performed best, likely reflecting the smaller number of available genomes. As filtering reduced the “All” reference set from 115,850 genomes across 1,020 lineages (global) to 1,678 genomes over 54 lineages (state), a higher similarity threshold was needed to maintain diversity among retained references. Thus, filtering by geographic proximity helps exclude unlikely lineages while focusing on variation among locally relevant lineages.

These results demonstrate that incorporating context-dependent metadata such as sampling location can complement purely sequence-based reference selection, leading to improved taxonomic profiling accuracy in geographically constrained viral datasets.

### Profiling accuracy improves with reference set size at species-level, but not below

In the bacterial species-level experiments we observe that larger reference sets generally result in higher profiling accuracy. In contrast, for the bacterial strain-level and viral experiments the reference sets with fewer genomes performed best. To investigate this relationship, Figure 5 shows the taxonomic profiling metrics (abundance accuracy and F1-score) versus the number of included reference genomes, for all profiling tools in the bacterial experiments. For the species-level experiments, both metrics correlate strongly with the number of genomes included (Spearman correlations between 0.875 and 0.966). This is also confirmed by the family- and order-based experiments (Figures S17–18 in Supplementary Material). For the strain-level experiments, however, we do not see such correlation, except for F1-score with Centrifuge (*ρ* = 0.649; Figure 5 panels D and E).

**Fig. 5.**
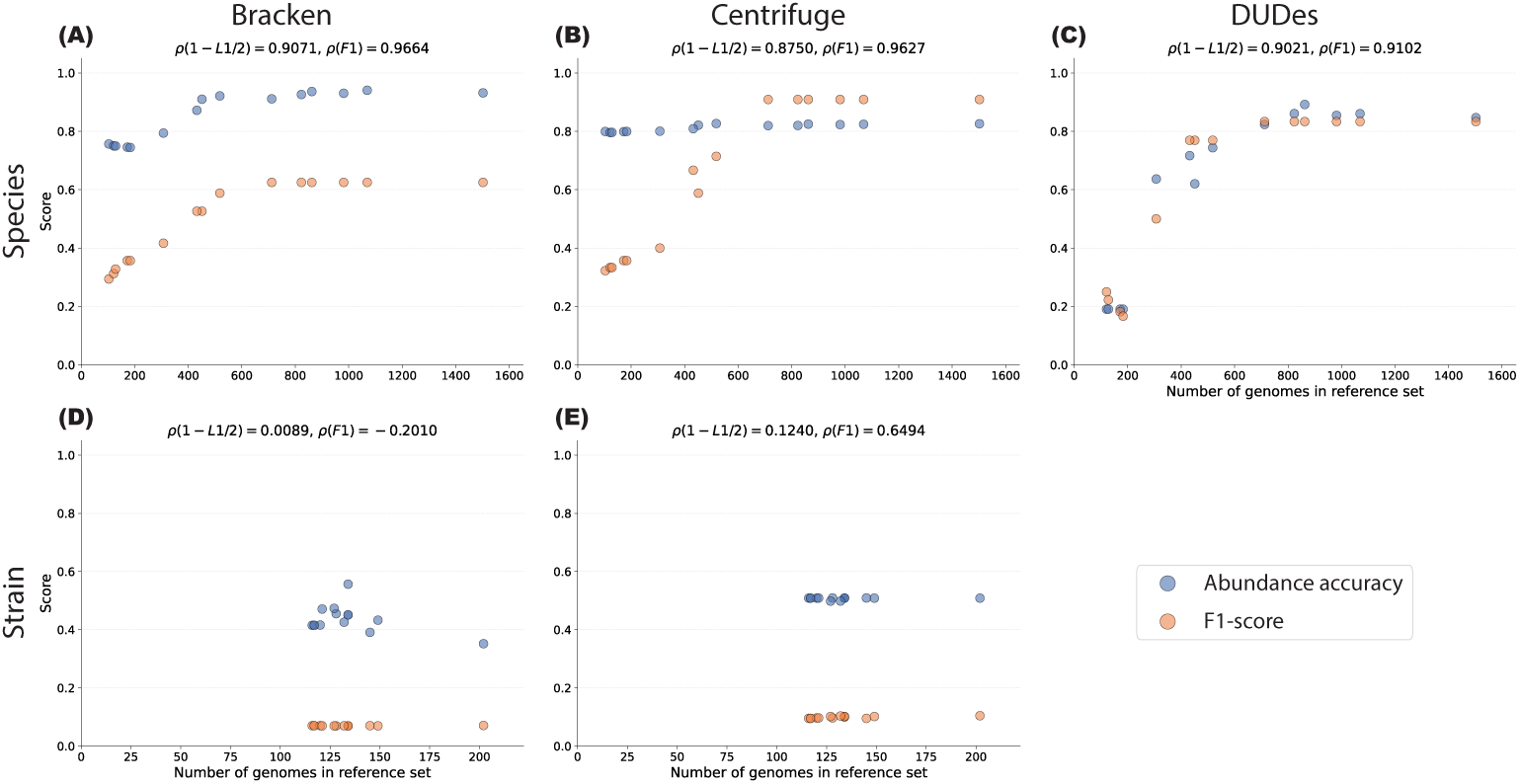
Median accuracy (abundance accuracy, F1-score) of taxonomic profiling tools across the bacterial experiments against the number of genomes in a reference set. Panels A-E are arranged in two rows (experimental setting: species-level, strain-level) and three columns (profiling tools: Bracken, Centrifuge, DUDes). *ρ*-values in subtitles show Spearman correlation between number of genomes included and median accuracy score.

For the viral experiments, we observe negative correlations: the F1-score decreases with increasing reference set sizes for both the global and country experiments (Figure 6**(A)-(B)**). Additionally, the correlation between reference set size and abundance accuracy is only 0.10 and 0.25 for the global and country experiments, respectively. In the state experiments (Figure 6(C)), there is an initial improvement in accuracy when reference sets grow, but this quickly plateaus. When aggregating results across all viral experiments, the Spearman correlations were negative (−0.57 for abundance accuracy and −0.73 for F1-score, respectively). Since the same samples were used in all viral experiments, this shows that smaller and more specific viral reference sets tend to achieve higher accuracies.

**Fig. 6.**
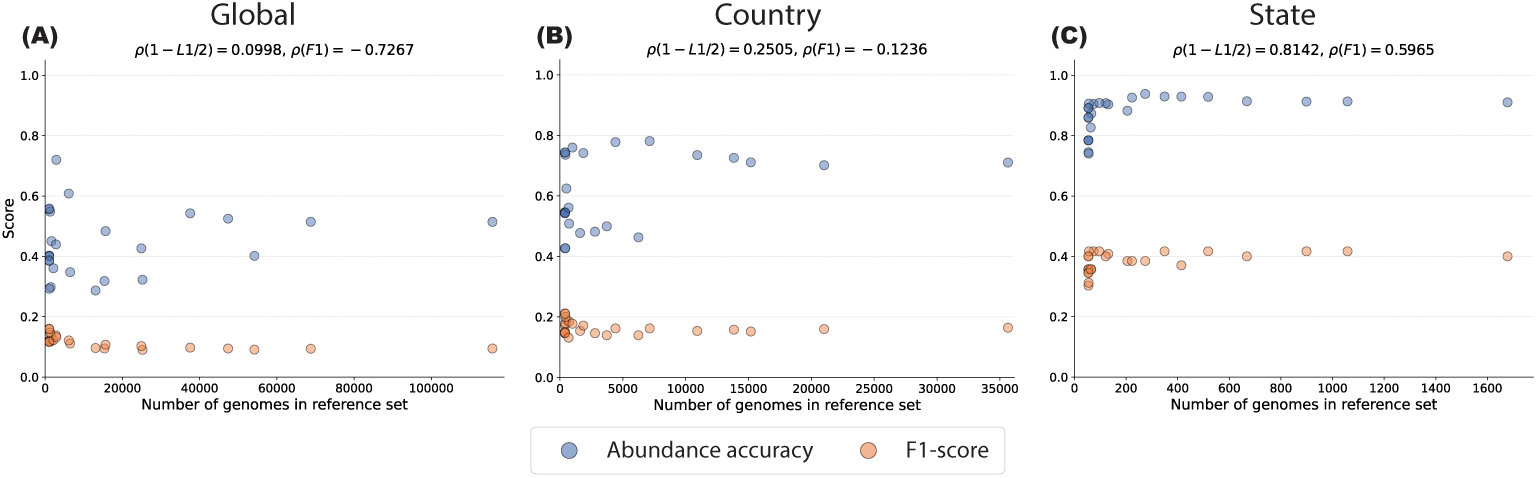
Median accuracy (abundance accuracy, F1-score) across SARS-CoV-2 experiments against the number of genomes in a reference set. Panels A–C correspond to the three geographic filtering levels: **(A)** no filter, **(B)** country filter (USA), and **(C)** state filter (Connecticut). *ρ*-values in subtitles show Spearman correlation between number of genomes included and median accuracy score.

Together, these results support the conclusion that the optimal reference set selection strategy depends on the particular context and profiling resolution of experiments.

### Validation on a real mock community confirms simulated results

To evaluate whether the improvements observed in simulated datasets translate to real sequencing data, we profiled the *E. coli* mock community sample presented in [14] after filtering host reads. Although derived from real sequencing data, this mock community has known strain abundances (see Methods), so that we can evaluate abundance accuracy and F1-score.

Figure 7 shows that performance metrics of all reference sets in the real mock community sample closely resemble those in the simulated analogue samples. In particular, several reference sets significantly improve abundance accuracy compared to the “All” reference set, while F1-scores remain similar, or slightly decrease. These results confirm that the benefits of reference genome selection for strain-level profiling generalize beyond simulated data.

**Fig. 7.**
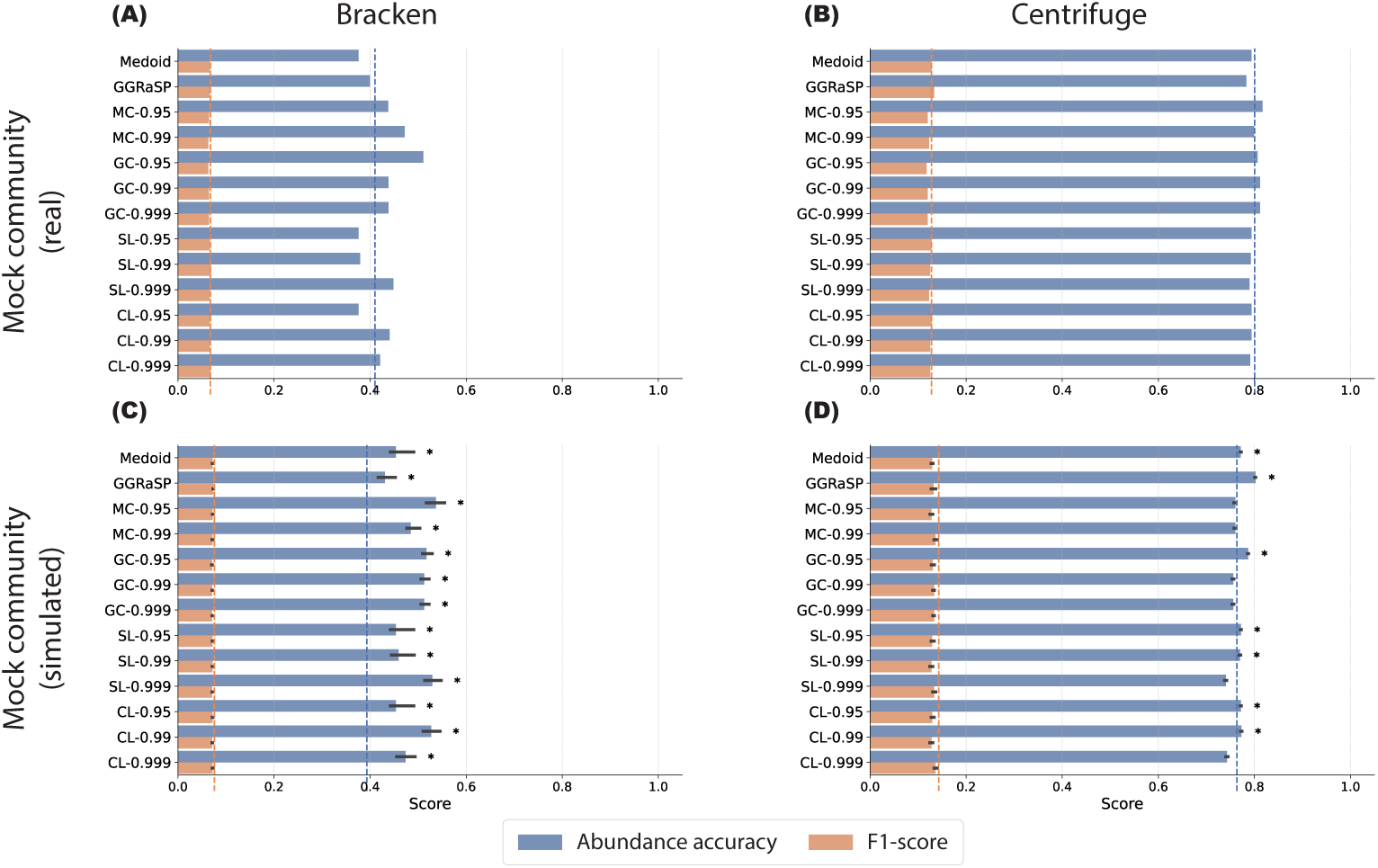
Accuracy metrics (abundance accuracy, F1-score) of taxonomic profiling tools on the mock community samples in the bacterial strain-level experiments. Panels A-D are arranged in two rows (sample type: real, simulated) and two columns (profiling tools: Bracken, Centrifuge). Dashed vertical lines correspond to the median performance of the baseline “All” reference set (no selection) and error bars correspond to the 10-90th percentile values. An asterisk indicates that the accuracy metric of a method is significantly better than that of the “All” reference set (Wilcoxon signed rank test, *p <* 0.05, Benjamini-Hochberg corrected).

While outcomes appear to differ between the real and simulated samples, with accuracy in the simulated samples generally being higher for most methods, these differences are not statistically significant (see Supplementary Table S11 in Supplementary Material). Existing discrepancies are likely the consequence of the additional processing steps of the real reads (primarily de-hosting), and to experimental noise that is absent in the simulated samples. Nevertheless, the overall trends remain consistent, supporting the conclusion that targeted reference genome selection can improve abundance estimation accuracy in realistic settings.

### Computational resource usage analysis

In addition to taxonomic profiling accuracy, another consideration when evaluating reference genome selection methods is their computational resource usage. To assess this, we measured total CPU time (system time + user time) and peak memory usage for all major processing steps (selection, indexing, and profiling). These measurements quantify the trade-off between the upfront cost of genome selection and the potential savings during indexing and repeated profiling. Note that dereplication tools can be run independently for each taxon, hence their effective runtime can be reduced through parallelization.

Figure 8 shows the runtimes for selection and indexing (combined) and taxonomic profiling across bacterial experiments, grouped by dereplication strategy: hierarchical clustering, MeShClust, Gclust, and All (no selection). Although indexing and profiling times differ between profilers, they show the same trends: hierarchical clustering methods generally result in lower overall runtime and memory usage during selection and indexing compared to the “All” selection, whereas MeShClust and Gclust show higher resource usage (see Supplementary Tables S2–3 for average resources per run). During profiling, runtimes for Bracken and Centrifuge were highly similar, regardless of index size (Figure 8 C,D). In contrast, profiling times for DUDes increased with index size, with runtimes exceeding four hours for the “All” reference set in the species-level experiments, and just under one hour for the medoid reference set.

**Fig. 8.**
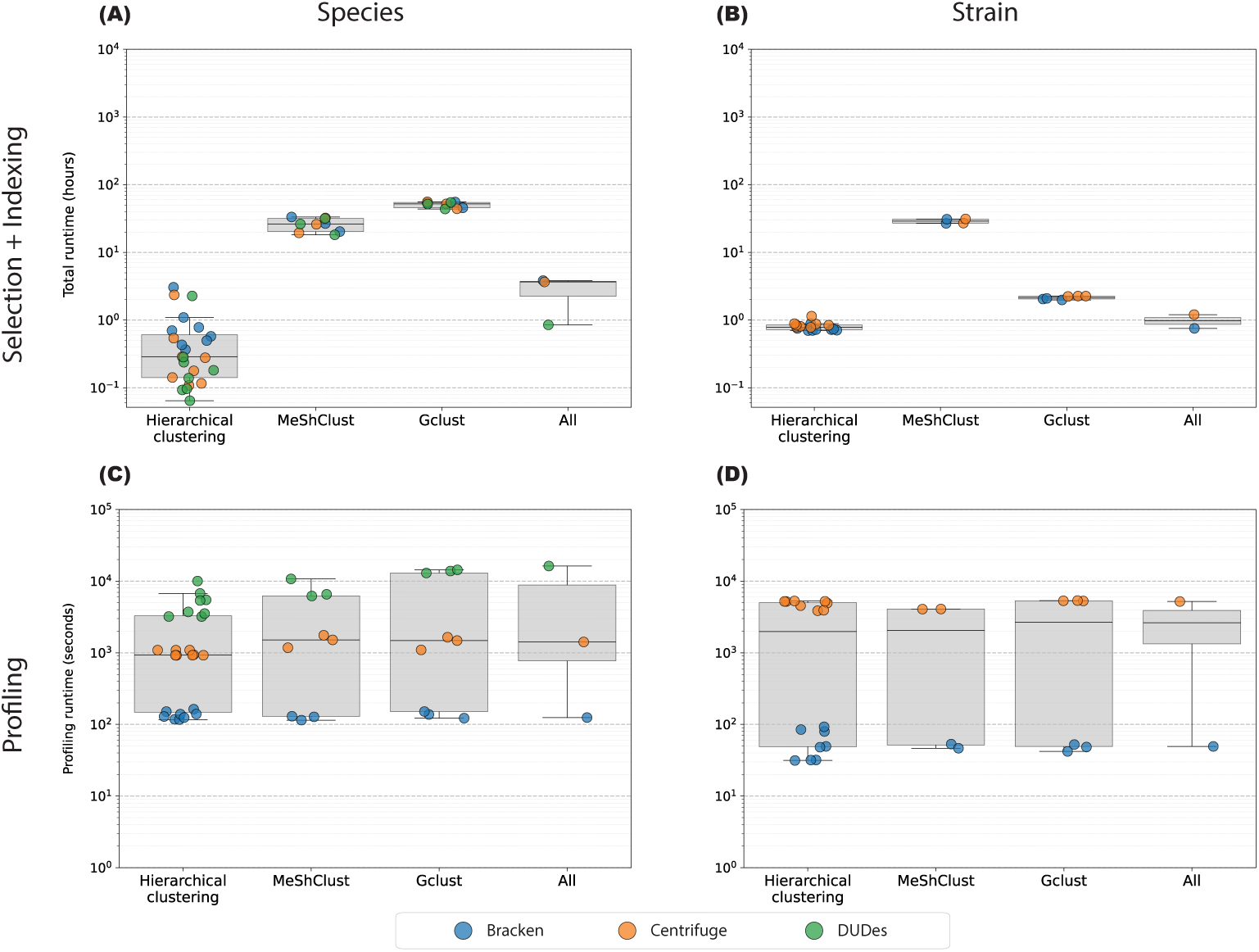
Runtime in bacterial experiments for reference selection, indexing, and taxonomic profiling. Dereplication methods are grouped by strategy: hierarchical clustering, MeShClust, Gclust, and All (no selection). Panels A&B show the average combined runtime for selection and indexing, and panels C&D show the average profiling runtime. Columns correspond to the experimental setting (species-level, strain-level). Profiling tools are color-coded: Bracken (blue), Centrifuge (orange), and DUDes (green). Whiskers in boxplots indicate 25–75th percentile values.

Although peak memory scales with index size for all profilers, this effect was modest for both Bracken and Centrifuge, with peak memory usage ranging between 0.3 GB and 1.4 GB in both the strain-level and species-level experiments (Supplementary Tables S2–3). For DUDes, the relationship was again more pronounced, requiring between 2.65 GB (medoid) and 26.47 GB (“All”) memory in the species-level experiments. The family- and order-based species-level experiments show that at larger scale (more taxa) the impact of reference selection on computational resource usage becomes stronger (Supplementary Tables S12–13).

In the viral experiments (Figure 9), indexing resource usage strongly scaled with reference set size, with the “All” reference set requiring the most time and memory (Supplementary Tables S4–6). However, when including the selection stage, other reference sets took longer overall. Peak memory usage scaled with reference set size, except for the VLQ-based selection which had a roughly constant memory footprint across experiments, resulting in relatively higher memory usage in the smaller state-filtered datasets. Profiling runtimes were determined primarily by index size, with the “All” selection consistently being the most resource-intensive.

**Fig. 9.**
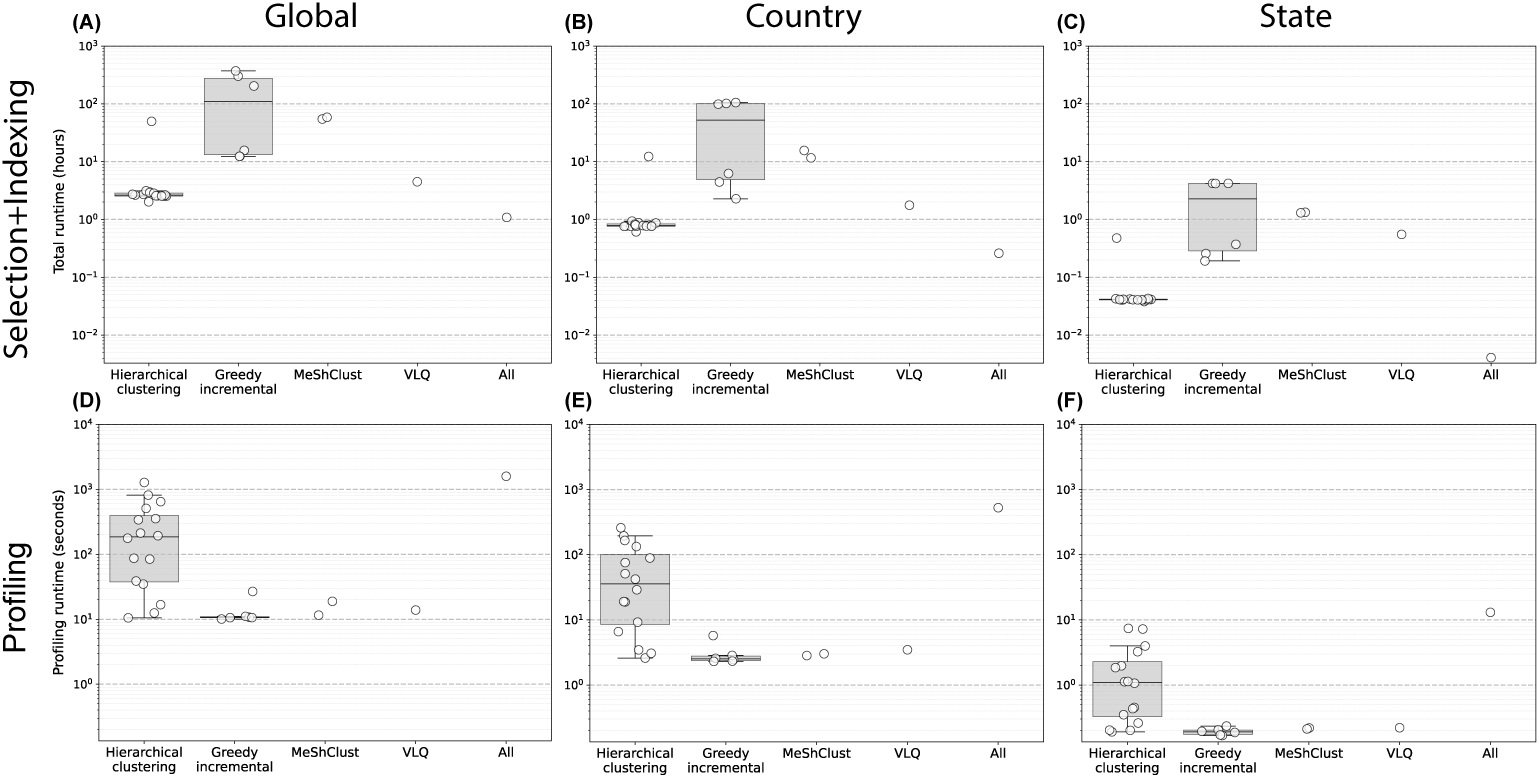
Runtime in SARS-CoV-2 experiments for reference selection, indexing, and taxonomic profiling. Dereplication methods are grouped by strategy: hierarchical clustering, MeShClust, Gclust, VLQ, and All (no selection). Panels A–C show the average combined runtime for selection and indexing, and panels D–F show the average profiling runtime. Columns correspond to the experimental setting (no filter, country filter, state filter). Whiskers in boxplots indicate 25–75th percentile values.

Overall, reference selection primarily benefits memory efficiency in bacterial profiling, while in the viral setting, both indexing and profiling times improved substantially due to the smaller number of reference genomes. This highlights that the computational benefits of reference genome selection depend on dataset size, genomic redundancy, and profiling method.

## Discussion

As genome databases continue to grow, identifying representative reference genomes has become essential to ensure computationally efficient downstream analyses [57]. Here, we systematically evaluated sequence dereplication methods for selecting representative viral and bacterial reference genomes, and quantified their impact on taxonomic profiling accuracy and computational performance. We estimated species- and strain-level abundances for bacterial datasets, and lineage abundances for viral datasets, finding that reference genome selection can substantially improve profiling accuracy at strain- and lineage-level, while also improving computational efficiency in the viral experiments. In contrast, for species-level bacterial profiling, reference genome selection yielded marginally higher accuracy with only a minor reduction in profiling runtime.

We believe that the gap between the effects observed is driven by fundamental differences in genomic diversity and redundancy, and taxonomic resolution. In the viral and strain-level bacterial experiments, abundance estimation was performed at higher resolution, which is intrinsically harder than species-level profiling. Although our viral experiments were limited to SARS-CoV-2 lineages, we expect similar patterns for other viruses: the impact of reference genome selection will depend largely on the level of taxonomic resolution being targeted. At finer resolutions, such as lineage or strain-level analyses, careful reference selection is likely to improve discrimination between closely related genomes. At coarser classification levels, such as the subtype structure of influenza A virus (e.g. H1N1, H3N2) or the four serotypes of dengue virus (DENV-1–4), the greater divergence between groups should reduce ambiguity in read assignment. As a result, abundance estimates are likely to be less sensitive to the choice of reference genomes.

Beyond sequence similarity-based dereplication, our viral experiments emphasize the utility of incorporating contextual metadata in reference genome selection. Previous work has shown that including temporal metadata or measures related to completeness can aid in selecting representative sequences [34, 40]. Our results extend this notion, showing that incorporating geographical proximity to samples further improves profiling accuracy, as SARS-CoV-2 lineages are often associated with specific regions. The applicability of such approaches will depend on the availability of contextual metadata associated with reference genomes. For example, repositories like GISAID routinely provide structured metadata including sampling date and geographic origin for a collection of pathogenic viruses [24]. However, it remains unclear to what extent this benefit generalizes to other viruses, or to bacterial datasets, where geographic signals may differ.

While several dereplication methods were shown to have a positive impact on taxonomic profiling accuracy, these methods depend strongly on a similarity threshold. For bacterial species-level profiling, higher thresholds consistently performed best in our experiments. In contrast, for higher resolution settings no universal threshold selection strategy emerged. This suggests that threshold selection should be treated as an integral and context-dependent component of reference genome selection for taxonomic profiling. Future work could focus on developing principled or adaptive approaches for selecting thresholds in similar high-resolution profiling tasks. In settings where such strategies are unavailable, auxiliary metrics such as the proportion of aligned reads can serve as a proxy indicator for evaluating thresholds, although this would require experimental validation.

Reference genome selection also introduces a practical trade-off between the upfront cost of selection and potential gains in downstream profiling efficiency and accuracy. Selection only marginally reduced profiling resource usage in the bacterial experiments, but in the viral setting it dramatically reduced profiling runtime and memory usage while simultaneously improving profiling accuracy. To reduce the upfront cost for reference selection, modern dereplication tools (e.g. LINCLUST [52] or CD-HIT [16]) accelerate large-scale clustering by filtering out clearly dissimilar sequence pairs. However, we expect these advantages to be less pronounced when applied to highly similar sequences, where many pairwise comparisons remain necessary.

Beyond selection costs, an inherent difficulty of performing experiments at this scale is the computational resource requirements for constructing large taxonomic profiling indexes [39]. We were therefore limited in scale, as constructing profiling indices using all available reference genomes (or other very large reference sets) would have required terabytes of memory and storage. It remains unclear whether our findings generalize across all bacterial datasets, highlighting the need for larger benchmarks, such as those in the CAMI challenges [38, 49], to more comprehensively assess the impact of reference genome selection on taxonomic profiling. Future work could also evaluate the interaction between reference genome selection and dedicated high-resolution profilers, such as StrainGE [14] and PanTax [59] for bacterial strain-level analysis, and VirStrain [30] and VirPool [17] for viral lineage profiling.

Given these limitations, a key next step is the development of reference selection methods designed specifically for taxonomic profiling. Although existing dereplication tools can be applied per taxon to ensure at least one genome is retained, this approach ignores inter-taxon similarities—an important factor for accurate classification. Developing methods that explicitly optimize discrimination between taxa could therefore improve both the accuracy and efficiency of taxonomic profiling, ultimately enabling more scalable and reliable analyses of microbial communities.

## Supplementary information

***Supplementary Material***

Includes Supplementary Information, all Supplementary Figures, and Supplementary Tables S1 and S7–11.

***Supplementary Table S2***

Computational resource usage for the bacterial strain-level experiments (including alignment statistics).

***Supplementary Table S3***

Computational resource usage for the genus-based bacterial species-level experiments (including alignment statistics).

***Supplementary Table S4***

Computational resource usage for the viral global experiments (including alignment statistics).

***Supplementary Table S5***

Computational resource usage for the viral country experiments (including alignment statistics).

***Supplementary Table S6***

Computational resource usage for the viral state experiments (including alignment statistics).

***Supplementary Table S12***

Computational resource usage for the family-based bacterial species-level experiments (including alignment statistics).

***Supplementary Table S13***

Computational resource usage for the order-based bacterial species-level experiments (including alignment statistics).

## List of abbreviations

bp: Basepairs
CI: Containment Index
GTDB: Genome Taxonomy Database
Mbp: Megabase pair
NCBI: National Center for Biotechnology Information
VLQ: Viral Lineage Quantification

## Declarations

### Ethics approval and consent to participate

Not applicable.

### Consent for publication

Not applicable.

### Availability of data and materials

The datasets supporting the conclusions of this article are available in a Zenodo repository [6] (10.5281/zenodo.18867106).

Supporting scripts as well as instructions for obtaining the results presented in this manuscript are available on Github [4].

- Project name: reference_set_selection_benchmark
- Project homepage: https://github.com/JaspervB-tud/reference_set_selectionbenchmark
- Archived version: 10.5281/zenodo.18888548
- Operating system(s): Linux
- Programming language: Python, Shell and R
- License: MIT

### Authors’ contributions

JvB, IN, and JAB conceived the study. IN designed and implemented the viral experiments. JvB implemented the reference selection strategies, designed and implemented the bacterial experiments, and performed the data analysis. JvB drafted the manuscript and JvB and JAB wrote the final version. All authors read and approved the manuscript.

## Acknowledgements

We gratefully acknowledge all data contributors, i.e., the Authors and their Originating laboratories responsible for obtaining the specimens, and their submitting laboratories for generating the genetic sequence and metadata and sharing via the GISAID Initiative, on which part of this research is based.

Research reported in this work was partially or completely facilitated by computational resources and support of the Delft AI Cluster (DAIC) at TU Delft (RRID: SCR 025091), but remains the sole responsibility of the authors, not the DAIC team.

