## Supplementary Material for "Benchmarking the impact of reference genome selection on taxonomic profiling accuracy"

March 10, 2026

#### Contents

**1 Selection of dereplication tools** **1**

**2 Selection of taxa for bacterial species-level experiments** **2**

**3 Read simulation set-up (Bacterial experiments)** **2**

**4 Statistical significance assessment of containment indices** **2**

**5 Bacterial reference set composition overlaps** **3**

**6 Core selections** **7**

**8 Accuracy versus reference set size** **14**

**9 Comparison between real and simulated mock community** **15**

### 1 Selection of dereplication tools

In all of our experiments, we wanted to include a comprehensive and representative range of dereplication tools, in order to cover available software implementations. We initially considered the following collection of tools:

- VSEARCH [1]
- Gclust [2]
- CD-HIT [3]
- MeShClust [4]
- GGRaSP [5]
- Hierarchical clustering (dRep-like [6])
- LINCLUST [7]

To assess the feasibility of running these tool at our intended scale (dereplication of 1,000–4,000 full genomes per taxon) we ran them (using default parameters and 32 threads where possible) on the SARS-CoV-2 genomes of the B.1.1.7 lineage (downsampled to 1,000 genomes as described in the main text), and the 4,725 *Escherichia coli* genomes. For GGRaSP and hierarchical clustering we estimated genomic distances using MASH [8], with $k = 21$ and $s = 1000$ for the *E. coli* genomes, and $k = 31$ and $s = 5000$ for the SARS-CoV-2 genomes. From this initial set of experiments, it turned out that CD-HIT was unable to produce a selection for both the SARS-CoV-2 and *E. coli* genomes within 5 days of runtime. LINCLUST failed to produce a selection for the SARS-CoV-2 genomes due to runtime errors. Thus, we were unable to include CD-HIT and LINCLUST in our evaluation.

#### 2 Selection of taxa for bacterial species-level experiments

For our bacterial species-level, we made a selection of taxa (genus, family, order) on which to run the selection tools and profiling pipelines, in order to keep computations feasible, but also consider the impact of reference genome selection with varying target similarity. The selected taxa were chosen to prioritize genomically diverse groups suitable for dereplication, and were obtained as follows: first, we filtered out any species with fewer than 100 genomes in the database, since our primary interest lies in dereplication. Afterwards, remaining species were sorted according to the average intra-species distance (estimated using MASH, $k = 21, \text{sketchsize} = 1000$), from highest to lowest to prioritize diversity. Taxa were then selected as the first taxon (at the corresponding level) to which 5 species belonged, by parsing the sorted species list in order. Although we selected 5 species from the selected taxa, all other species of the selected taxa with at least on "Complete Genome" quality assembly on NCBI Assembly (now NCBI Genomes) were also included during dereplication and profiling. The selected taxa and species for all species-level experiments can be found in Supplementary Table S1 below.

Table S1: Experimental set-up: selected genus, family and order taxa, including the corresponding species selected for every experiment.

| Experiment | Selected taxon | Selected species |
|---|---|---|
| Genus | *Streptococcus* | *Streptococcus suis*<br>*Streptococcus thermophilus*<br>*Streptococcus agalactiae*<br>*Streptococcus pneumoniae*<br>*Streptococcus pyogenes* |
| Family | *Enterobacteriaceae* | *Citrobacter freundii*<br>*Escherichia coli*<br>*Salmonella enterica*<br>*Enterobacter hormaechei*<br>*Klebsiella quasipneumoniae* |
| Order | Enterobacterales | *Buchnera aphidicola*<br>*Citrobacter freundii*<br>*Escherichia coli*<br>*Serratia marcescens*<br>*Enterobacter hormaechei* |

#### 3 Read simulation set-up (Bacterial experiments)

For both the species- and strain-level bacterial experiments, we downloaded assemblies from NCBI Assembly to simulate reads from. For the species-level experiments, we used the top 10 assemblies (based on completeness), which were required to have a "Scaffold" or "Chromosome" assembly quality in order to prevent overlap between reference sets and simulation genomes.

For the strain-level experiments, we sorted assemblies on N50 (descending), selecting the highest ranked assembly not included in any reference where possible. An exception to this is the Sakai strain, for which only a single assembly was available, which was also included in the reference sets. Across all experiments, species or strain abundances were fixed at equal abundance ($\sim 20\%$ for species-level, $\sim 25\%$ for strain-level), uniformly distributed over all assemblies, with taxonomic abundances varying based on genome lengths.

#### 4 Statistical significance assessment of containment indices

Throughout our experiments, we calculate containment indices (CIs) between reference sets, to assess their similarities. The containment index between two reference sets $R_1, R_2$ is defined as $\text{CI}(R_1, R_2) = \frac{|R_1 \cap R_2|}{|R_1|}$. In this we only consider genomes from taxa that have at least 2 sequences, as otherwise the intersection for a given taxon is by definition equal to 1.

To assess the statistical significance of a containment index value, we constructed an empirical null distribution using Monte Carlo simulations. Specifically, for every taxonomic group $g$ (species for bacterial species-level experiments, strains for bacterial strain-level experiments and lineages for viral experiments), given the total number of genomes available $N_g$ and observed counts $n_{1g}$ and $n_{2g}$ in the reference sets $R_1$ and $R_2$ respectively, we sample overlaps from a hypergeometric distribution to simulate the random selection of $n_{1g}$ genomes, given $n_{2g}$ genomes already chosen. Summing simulated overlaps across all taxonomic groups yields a total overlap per simulation. Afterwards, we compute the corresponding containment index for every simulated draw (10,000 per containment index), and compare against the observed containment index to obtain a one-sided p-value testing whether the observed containment index was larger than expected.

#### 5 Bacterial reference set composition overlaps

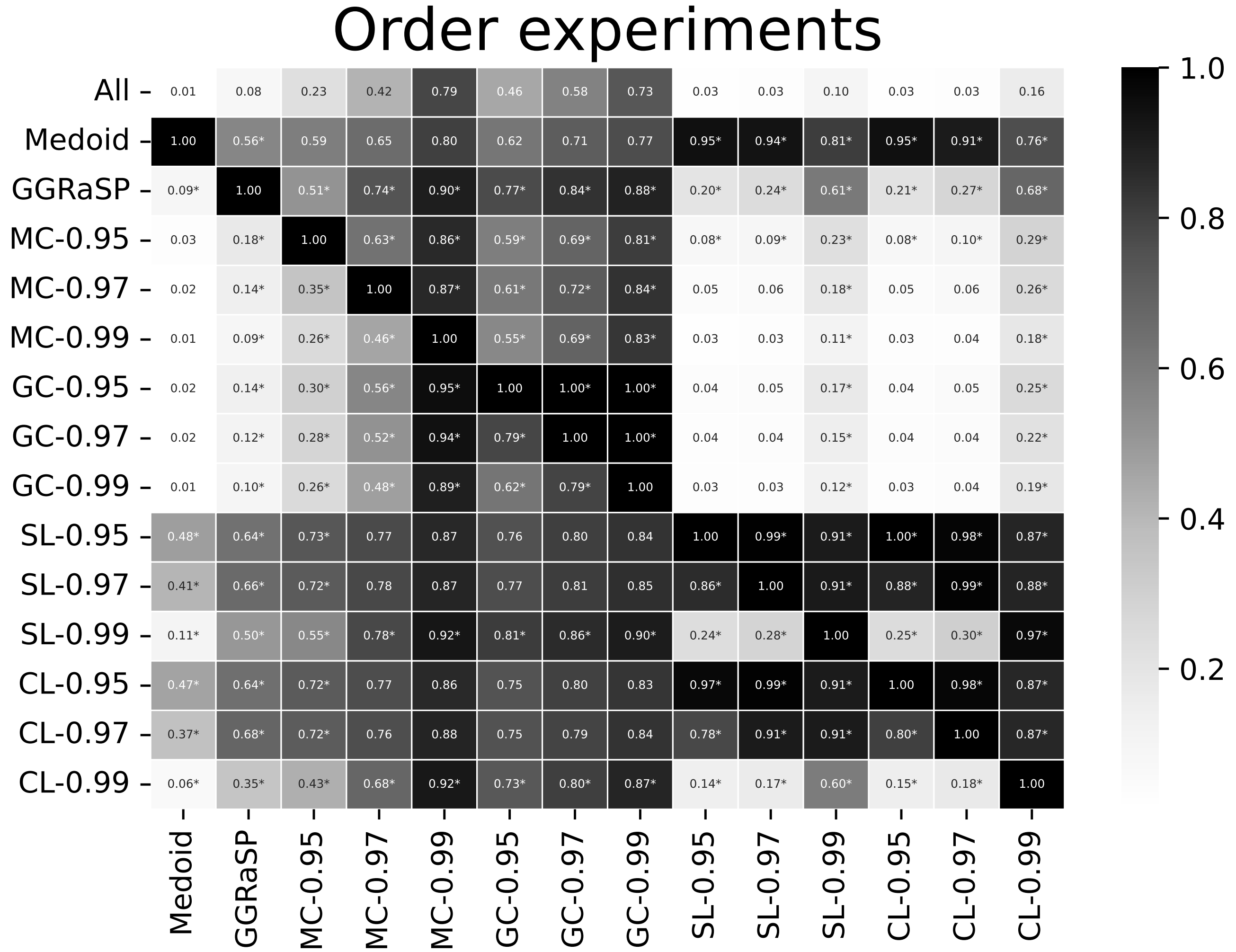

Figure S1: Containment indices (CIs) between reference genomes selected by different dereplication tools for the order-based species-level bacterial experiments. CI is only calculated for species that have more than 1 genome to select from. Asterisks indicate that the CI is significantly higher compared to randomly selecting genomes ($p < 0.05$, Benjamini-Hochberg corrected).

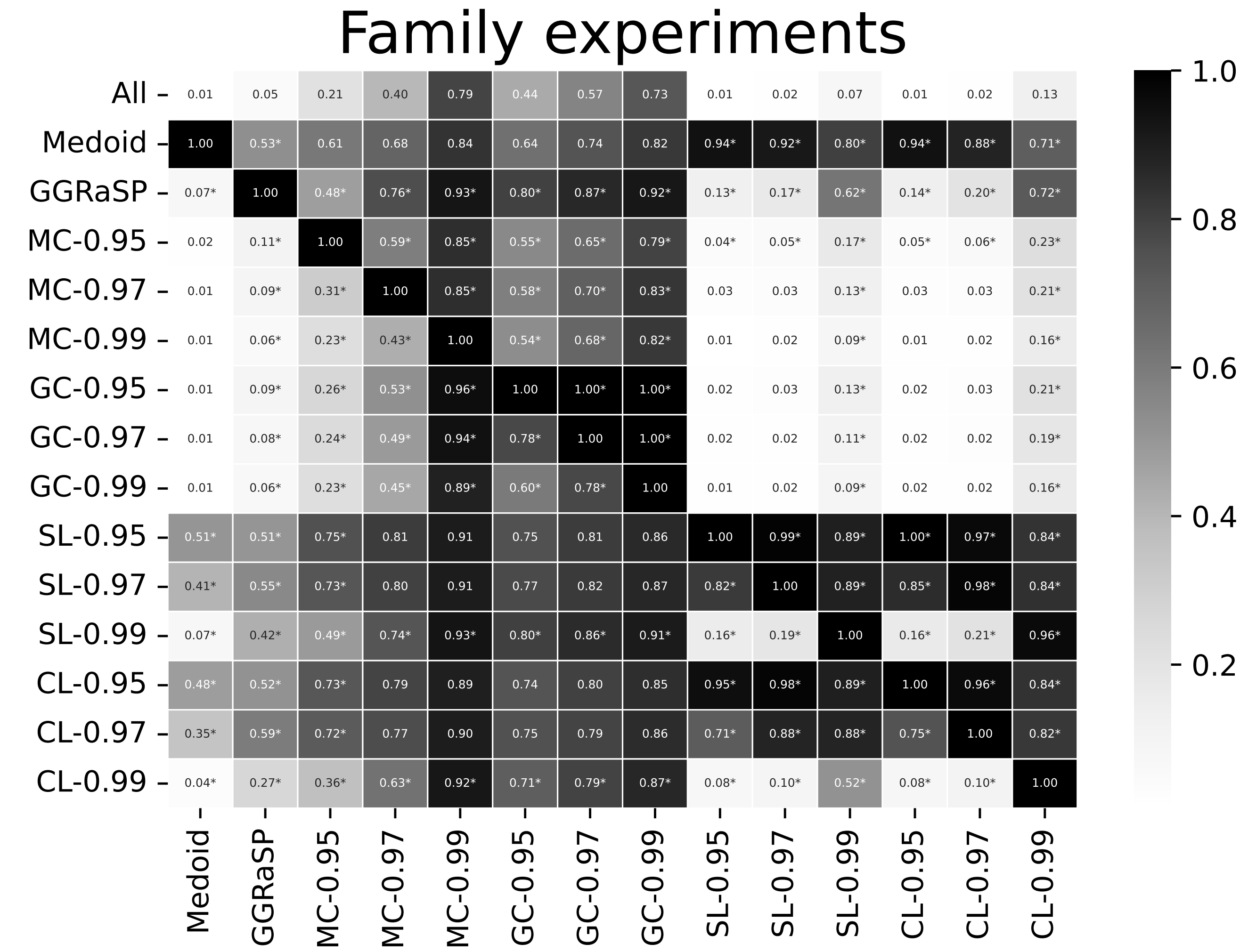

Figure S2: Containment indices (CIs) between reference genomes selected by different dereplication tools for the family-based species-level bacterial experiments. CI is only calculated for species that have more than 1 genome to select from. Asterisks indicate that the CI is significantly higher compared to randomly selecting genomes ($p < 0.05$, Benjamini-Hochberg corrected).

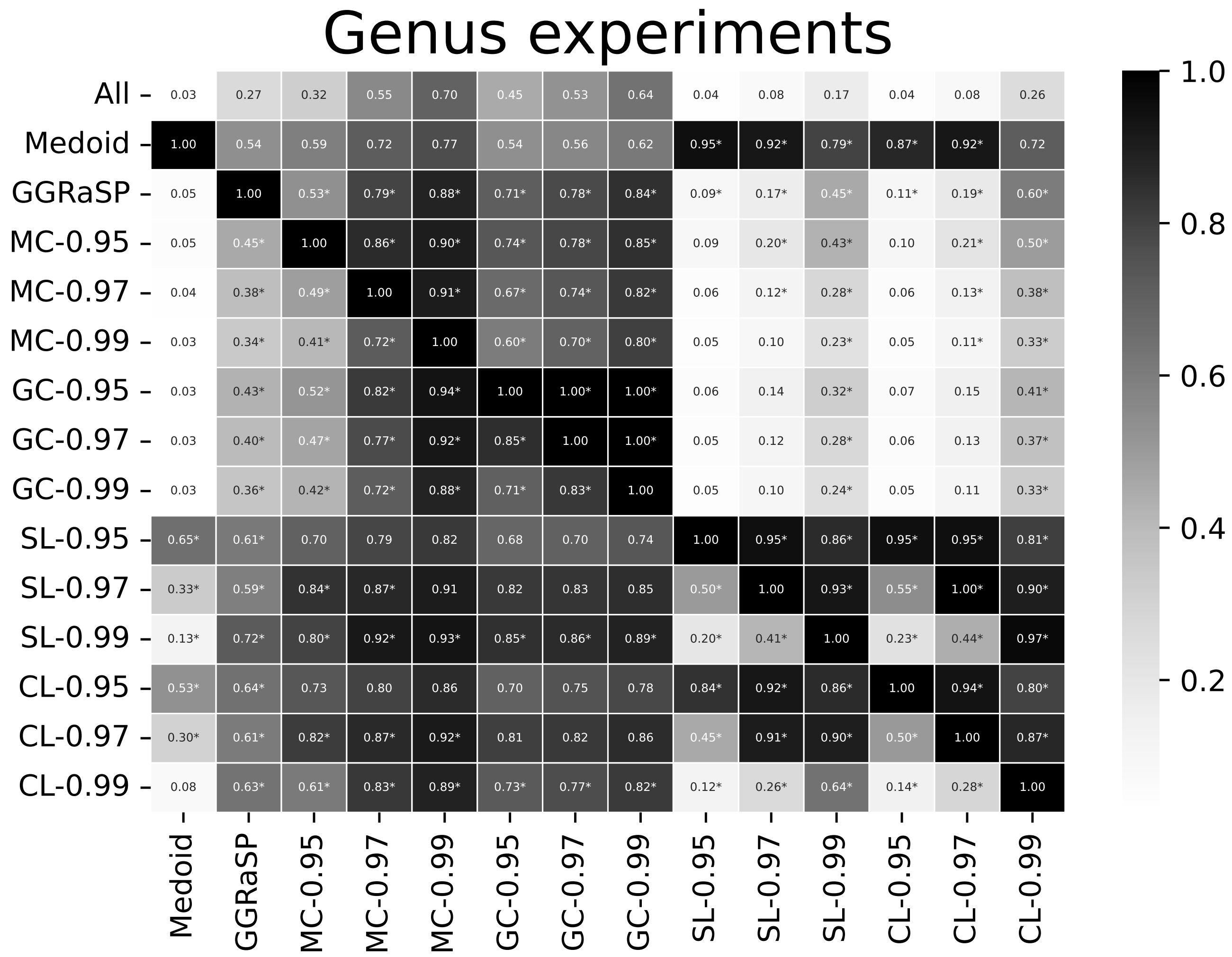

Figure S3: Containment indices (CIs) between reference genomes selected by different dereplication tools for the genus-based species-level bacterial experiments. CI is only calculated for species that have more than 1 genome to select from. Asterisks indicate that the CI is significantly higher compared to randomly selecting genomes ($p < 0.05$, Benjamini-Hochberg corrected).

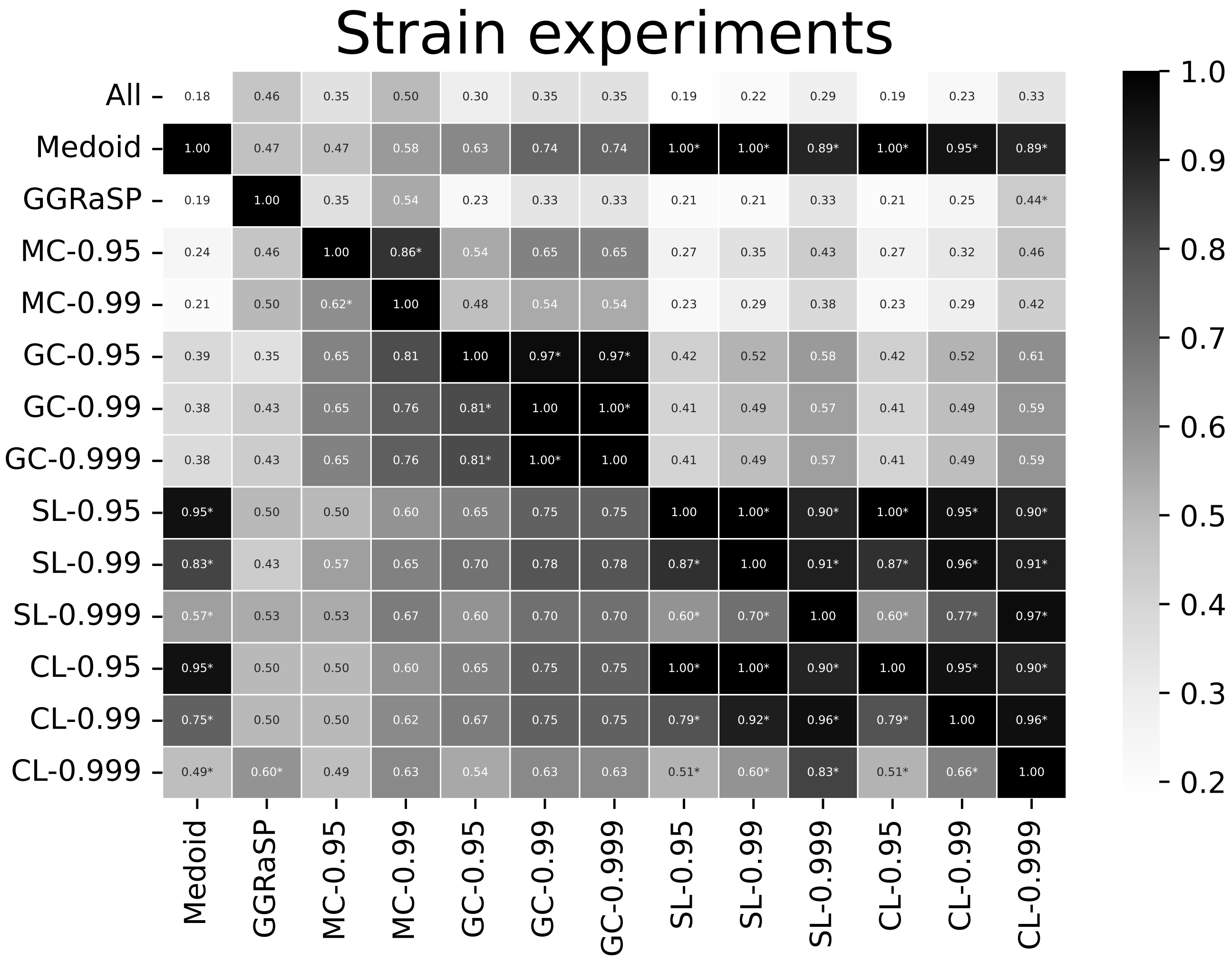

Figure S4: Containment indices (CIs) between reference genomes selected by different dereplication tools for the strain-level bacterial experiments. CI is only calculated for strains that have more than 1 genome to select from. Asterisks indicate that the CI is significantly higher compared to randomly selecting genomes ($p < 0.05$, Benjamini-Hochberg corrected).

### 6 Core selections

#### 6.1 Viral experiments

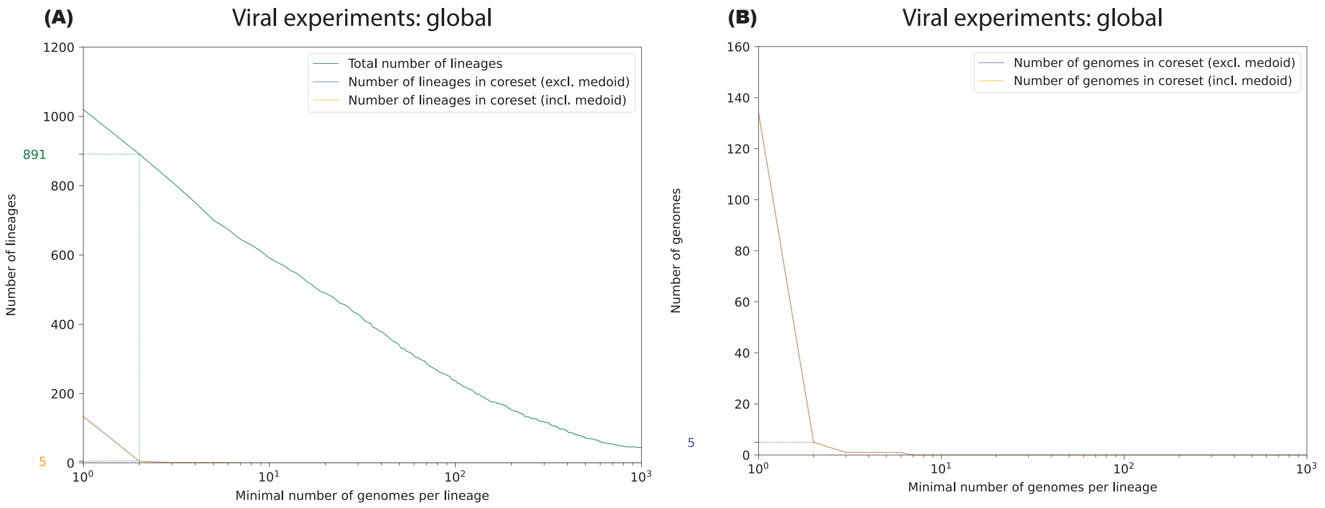

Figure S5: Coresets of reference genomes for the viral global experiments. **(A)** Number of lineages where every reference set shares at least 1 genome (y-axis) versus the lineages that have at least X genomes (x-axis). **(B)** Number of genomes selected by all reference sets (y-axis) versus the lineages that have at least X genomes (x-axis)

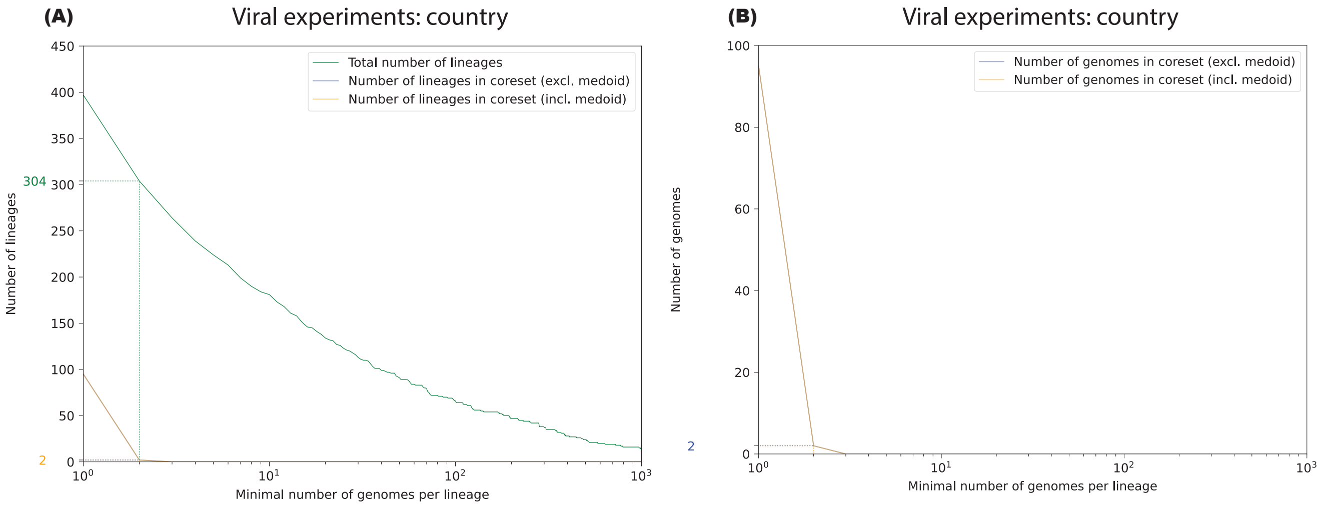

Figure S6: Coresets of reference genomes for the viral country experiments. **(A)** Number of lineages where every reference set shares at least 1 genome (y-axis) versus the lineages that have at least X genomes (x-axis). **(B)** Number of genomes selected by all reference sets (y-axis) versus the lineages that have at least X genomes (x-axis)

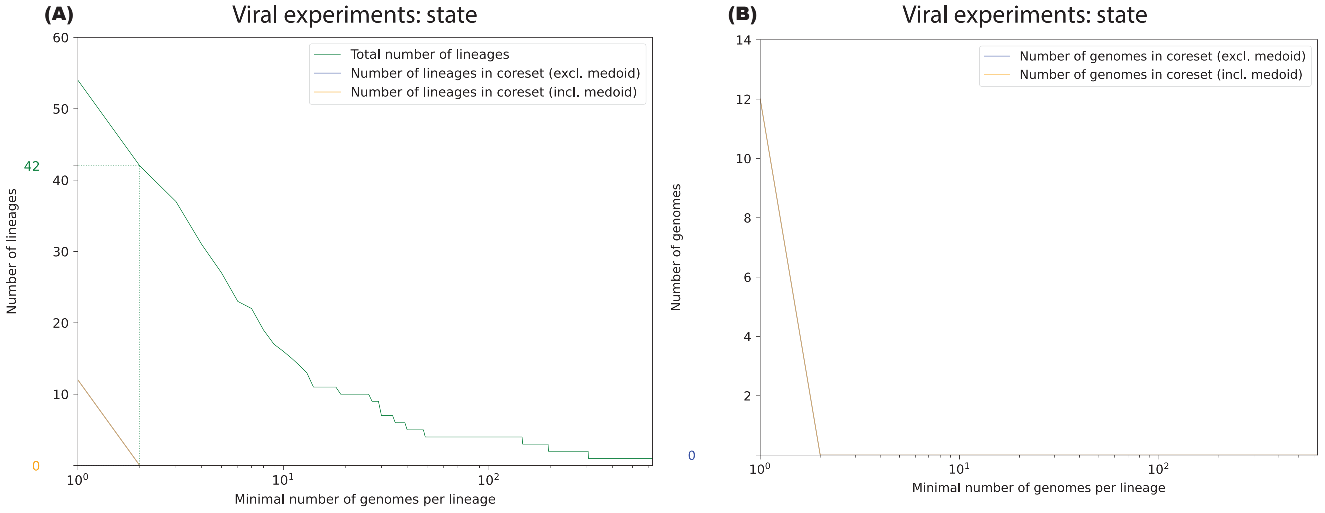

Figure S7: Coresets of reference genomes for the viral state experiments. **(A)** Number of lineages where every reference set shares at least 1 genome (y-axis) versus the lineages that have at least X genomes (x-axis). **(B)** Number of genomes selected by all reference sets (y-axis) versus the lineages that have at least X genomes (x-axis)

#### 6.2 Bacterial experiments

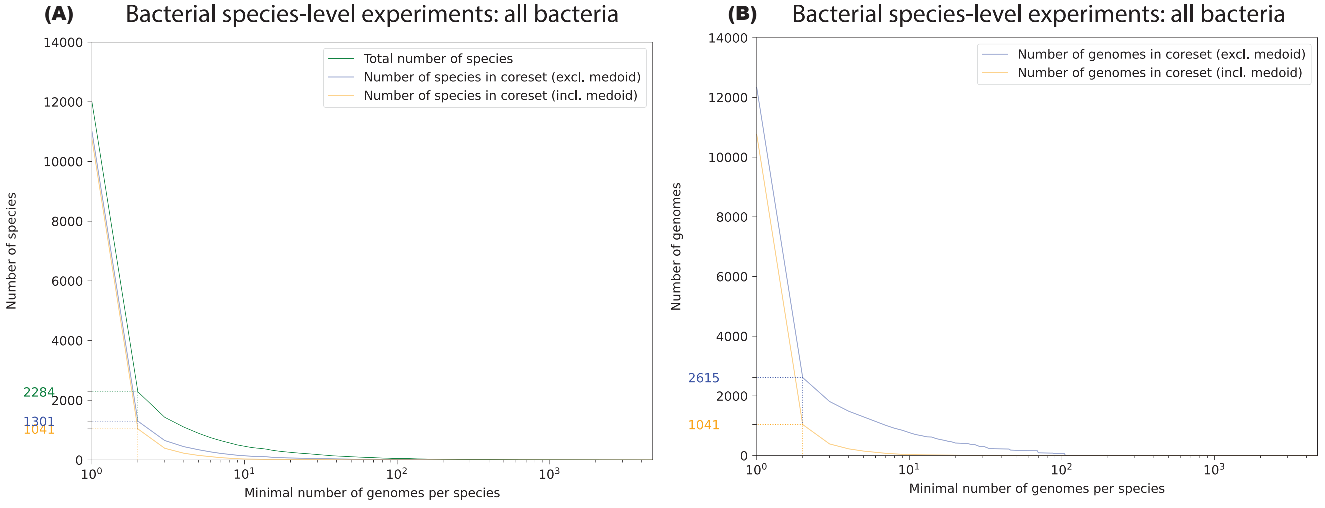

Figure S8: Coresets of reference genomes for the bacterial species-level all taxa experiments. **(A)** Number of species where every reference set shares at least 1 genome (y-axis) versus the species that have at least X genomes (x-axis). **(B)** Number of genomes selected by all reference sets (y-axis) versus the species that have at least X genomes (x-axis)

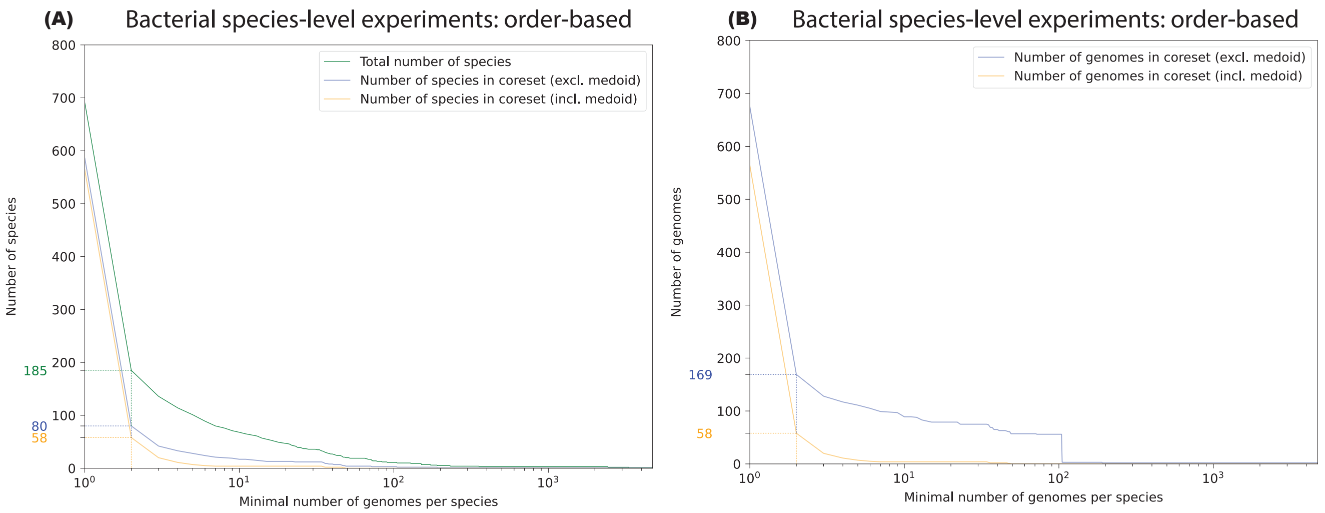

Figure S9: Coresets of reference genomes for the bacterial species-level order-based experiments. **(A)** Number of species where every reference set shares at least 1 genome (y-axis) versus the species that have at least X genomes (x-axis). **(B)** Number of genomes selected by all reference sets (y-axis) versus the species that have at least X genomes (x-axis)

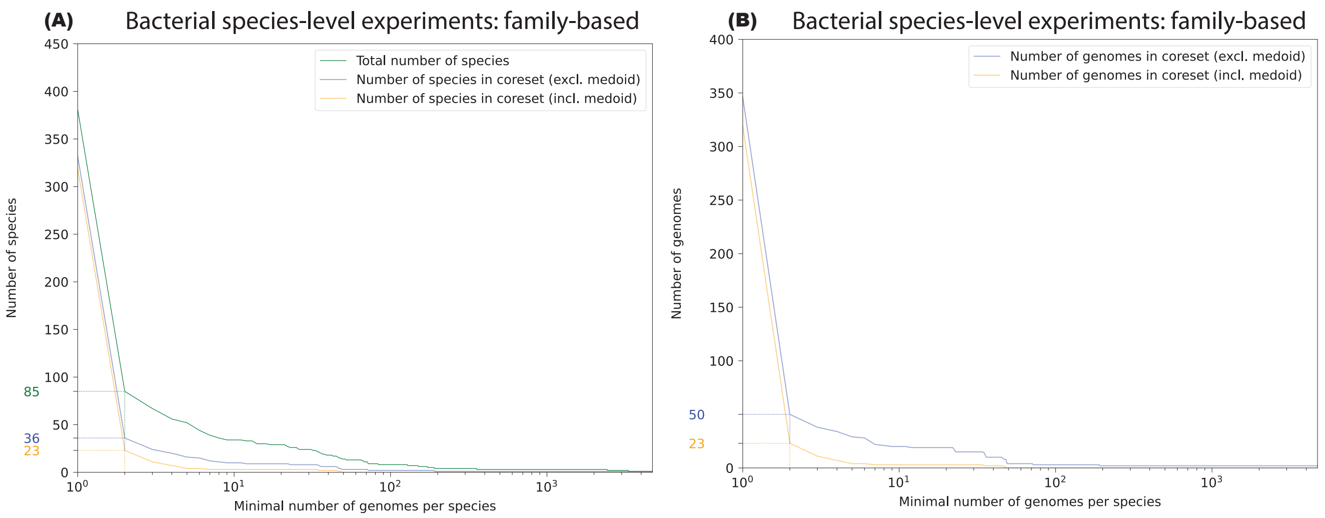

Figure S10: Coresets of reference genomes for the bacterial species-level family-based experiments. **(A)** Number of species where every reference set shares at least 1 genome (y-axis) versus the species that have at least X genomes (x-axis). **(B)** Number of genomes selected by all reference sets (y-axis) versus the species that have at least X genomes (x-axis)

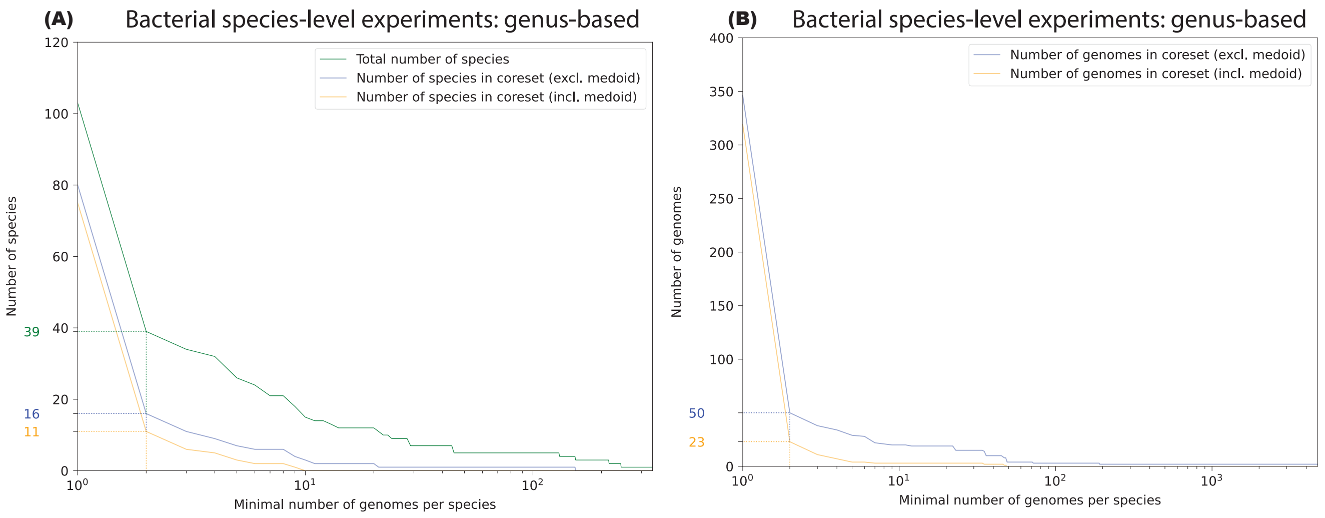

Figure S11: Coresets of reference genomes for the bacterial species-level genus-based experiments. **(A)** Number of species where every reference set shares at least 1 genome (y-axis) versus the species that have at least X genomes (x-axis). **(B)** Number of genomes selected by all reference sets (y-axis) versus the species that have at least X genomes (x-axis)

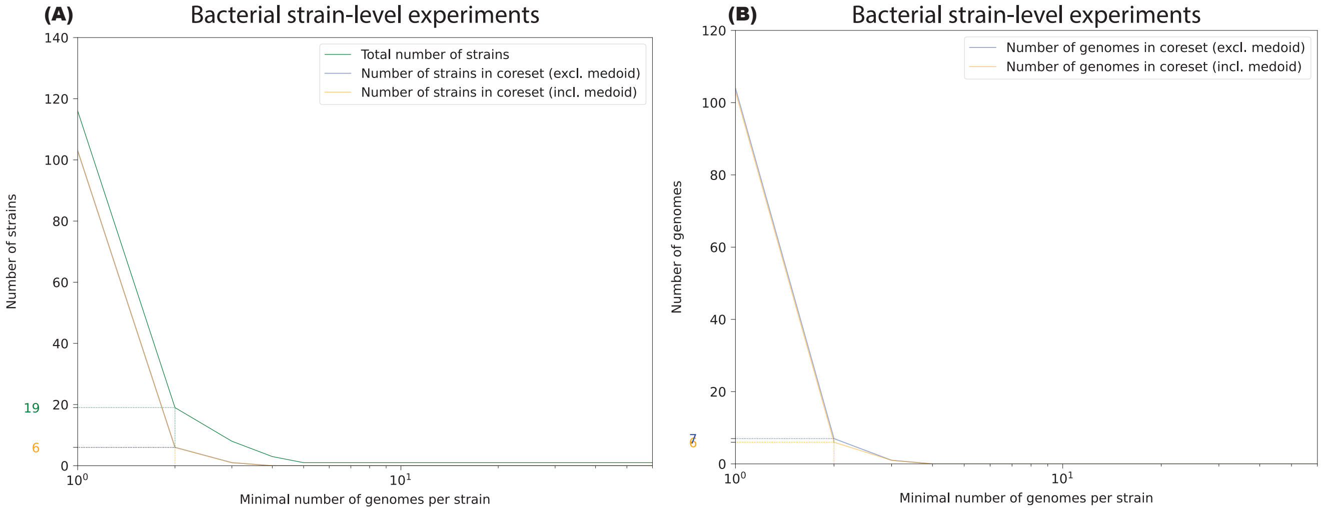

Figure S12: Coresets of reference genomes for the bacterial strain-level experiments. **(A)** Number of strains where every reference set shares at least 1 genome (y-axis) versus the strains that have at least X genomes (x-axis). **(B)** Number of genomes selected by all reference sets (y-axis) versus the strains that have at least X genomes (x-axis)

### 7 Profiling accuracy

#### 7.1 Bacterial experiments

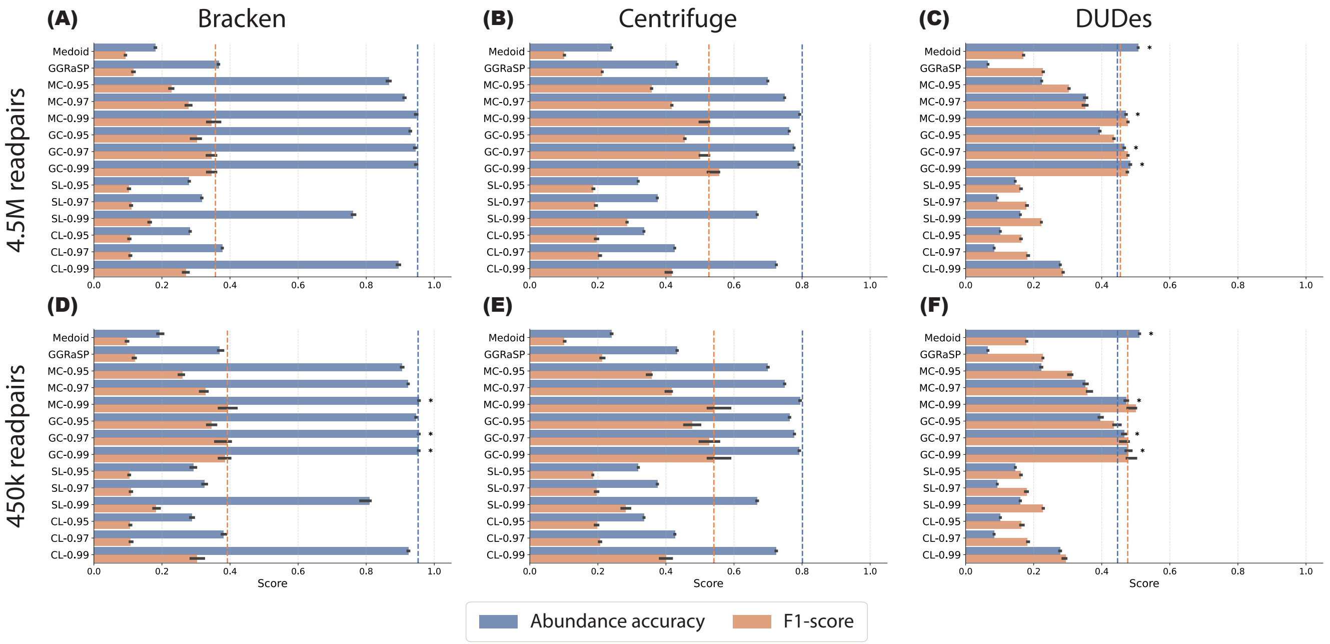

Figure S13: Accuracy metrics (abundance accuracy, F1-score) of taxonomic profiling tools in the bacterial order-based, species-level experiments. Panels A-F are arranged in two rows (readcount: 4.5 million readpairs, 450 thousand readpairs) and three columns (profiling tools: Bracken, Centrifuge, DUDes). Dashed vertical lines correspond to the median performance of the baseline "All" reference set (no selection) and error bars correspond to 10-90th percentile values. An asterisk indicates that the accuracy metric of a method is significantly better than that of the "All" reference set (Wilcoxon signed rank test, $p < 0.05$, Benjamini-Hochberg corrected).

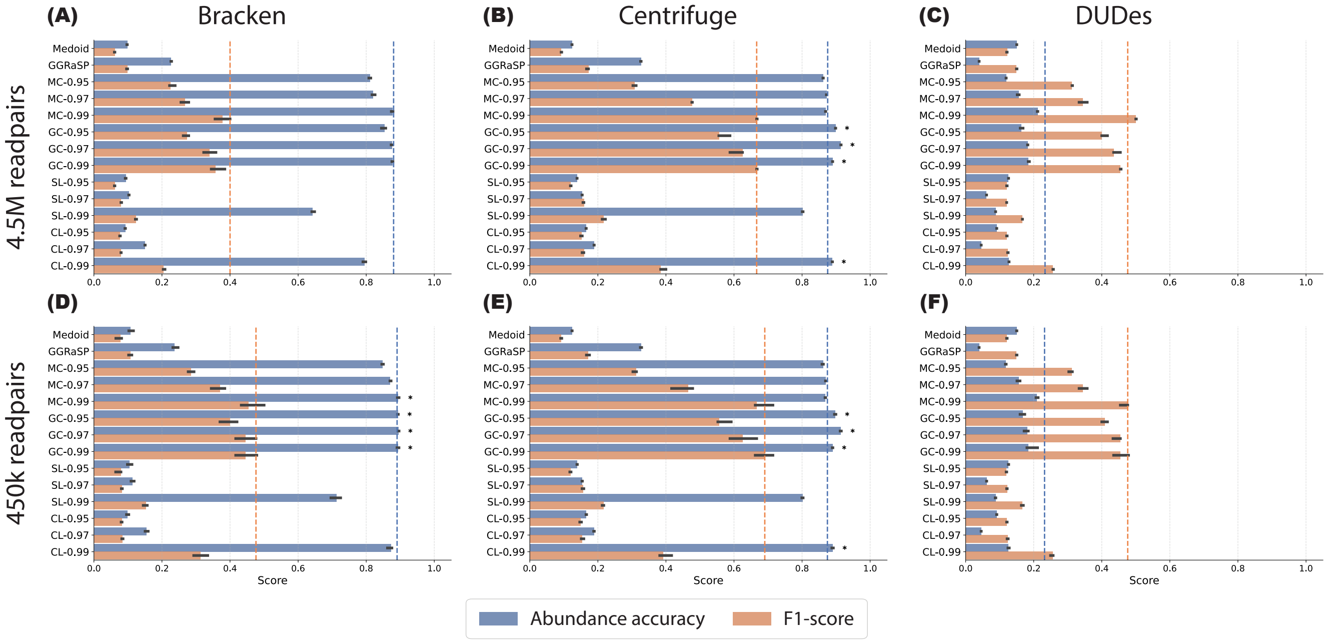

Figure S14: Accuracy metrics (abundance accuracy, F1-score) of taxonomic profiling tools in the bacterial family-based, species-level experiments. Panels A-F are arranged in two rows (readcount: 4.5 million readpairs, 450 thousand readpairs) and three columns (profiling tools: Bracken, Centrifuge, DUDes). Dashed vertical lines correspond to the median performance of the baseline "All" reference set (no selection) and error bars correspond to 10-90th percentile values. An asterisk indicates that the accuracy metric of a method is significantly better than that of the "All" reference set (Wilcoxon signed rank test, $p < 0.05$, Benjamini-Hochberg corrected).

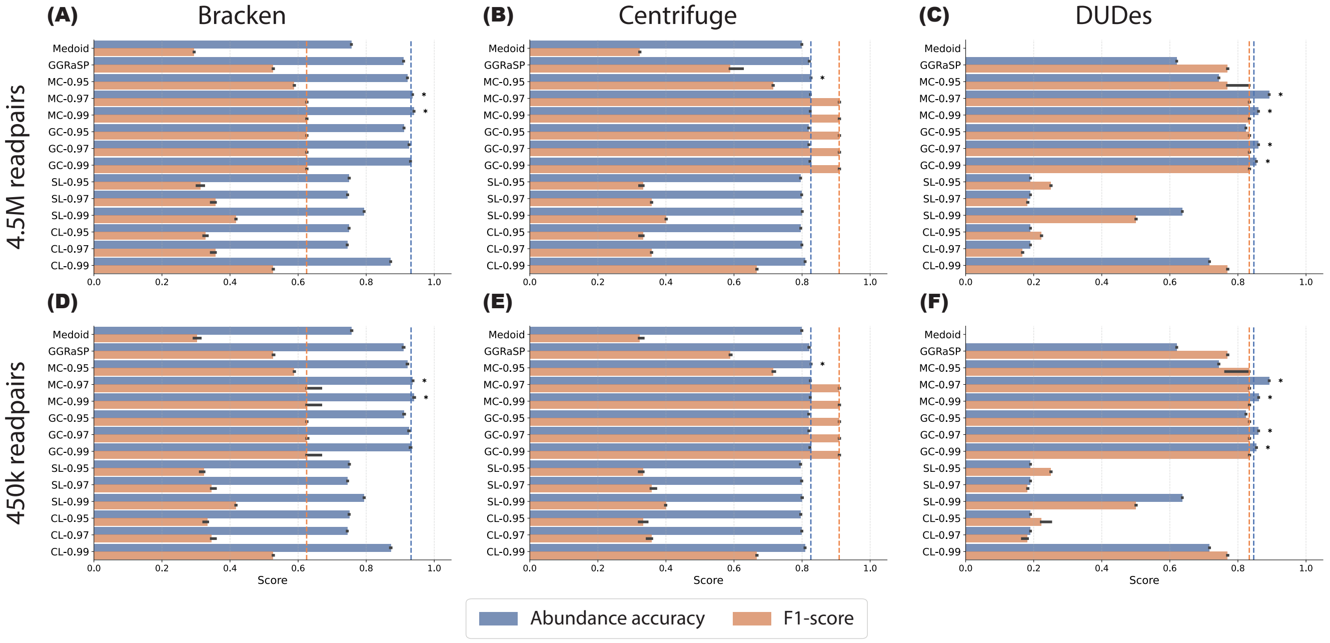

Figure S15: Accuracy metrics (abundance accuracy, F1-score) of taxonomic profiling tools in the bacterial genus-based, species-level experiments. Panels A-F are arranged in two rows (readcount: 4.5 million readpairs, 450 thousand readpairs) and three columns (profiling tools: Bracken, Centrifuge, DUDes). Dashed vertical lines correspond to the median performance of the baseline "All" reference set (no selection) and error bars correspond to 10-90th percentile values. An asterisk indicates that the accuracy metric of a method is significantly better than that of the "All" reference set (Wilcoxon signed rank test, $p < 0.05$, Benjamini-Hochberg corrected).

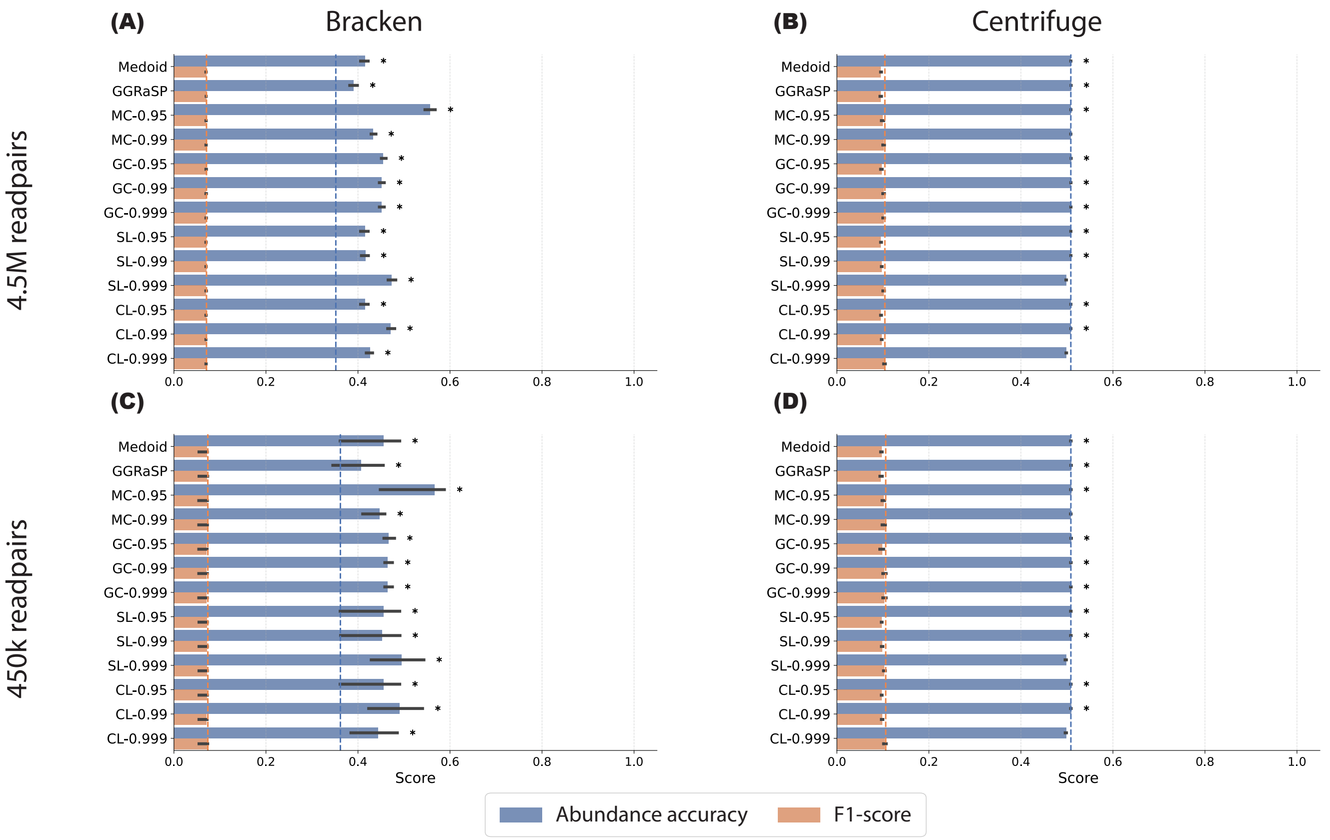

Figure S16: Accuracy metrics (abundance accuracy, F1-score) of taxonomic profiling tools in the bacterial strain-level experiments. Panels A-D are arranged in two rows (readcount: 4.5 million readpairs, 450 thousand readpairs) and two columns (profiling tools: Bracken, Centrifuge). Dashed vertical lines correspond to the median performance of the baseline "All" reference set (no selection) and error bars correspond to 10-90th percentile values. An asterisk indicates that the accuracy metric of a method is significantly better than that of the "All" reference set (Wilcoxon signed rank test, $p < 0.05$, Benjamini-Hochberg corrected).

**Testing for statistical significant differences based on readcount in bacterial experiments** In addition to providing the results for both readpair counts in all bacterial experiments, we have also assessed whether the difference is significant. More specifically, we calculate for every selection (excluding "All") and every sample, the difference in accuracy metric between the method and the "All" reference set outcome in both the 450k readpairs samples and 4.5m readpairs samples. To assess if our inference (namely the performance w.r.t. the "All" reference set) changes, we perform a two-sided Mann-Whitney U test between the 4.5m readpairs and 450k readpairs outcomes, correcting for multiple testing using the Benjamini-Hochberg correction.

Table S7: Assessment of statistical difference in outcome per accuracy metric (abundance accuracy, F1-score), per profiler, for bacterial order-based species-level experiment between 450 thousand readpairs and 5 million readpairs samples (two-sided Mann-Whitney U test, $p < 0.05$, Benjamini-Hochberg corrected). An asterisk indicates that there is a significant difference in scores between the two sequencing depths.

| | **Bracken** | | **Centrifuge** | | **DUDes** | |
|---|---|---|---|---|---|---|
| Method | Abundance accuracy corrected p-value | F1-score corrected p-value | Abundance accuracy corrected p-value | F1-score corrected p-value | Abundance accuracy corrected p-value | F1-score corrected p-value |
| Medoid | 1.000000 | 1.000000 | 1.000000 | 1.000000 | 1.000000 | 1.000000 |
| GGRaSP | 1.000000 | 1.000000 | 1.000000 | 1.000000 | 1.000000 | 1.000000 |
| MC-0.95 | 1.000000 | 1.000000 | 1.000000 | 1.000000 | 1.000000 | 1.000000 |
| MC-0.97 | 1.000000 | 1.000000 | 1.000000 | 1.000000 | 1.000000 | 1.000000 |
| MC-0.99 | 1.000000 | 1.000000 | 1.000000 | 1.000000 | 1.000000 | 1.000000 |
| GC-0.95 | 1.000000 | 1.000000 | 1.000000 | 1.000000 | 1.000000 | 1.000000 |
| GC-0.97 | 1.000000 | 1.000000 | 1.000000 | 1.000000 | 1.000000 | 1.000000 |
| GC-0.99 | 1.000000 | 1.000000 | 1.000000 | 1.000000 | 1.000000 | 1.000000 |
| SL-0.95 | 1.000000 | 1.000000 | 1.000000 | 1.000000 | 1.000000 | 1.000000 |
| SL-0.97 | 1.000000 | 1.000000 | 1.000000 | 1.000000 | 1.000000 | 1.000000 |
| SL-0.99 | 1.000000 | 1.000000 | 1.000000 | 1.000000 | 1.000000 | 1.000000 |
| CL-0.95 | 1.000000 | 1.000000 | 1.000000 | 1.000000 | 1.000000 | 1.000000 |
| CL-0.97 | 1.000000 | 1.000000 | 1.000000 | 1.000000 | 1.000000 | 1.000000 |
| CL-0.99 | 1.000000 | 1.000000 | 1.000000 | 1.000000 | 1.000000 | 1.000000 |

Table S8: Assessment of statistical difference in outcome per accuracy metric (abundance accuracy, F1-score), per profiler, for bacterial family-based species-level experiment between 450 thousand readpairs and 5 million readpairs samples (two-sided Mann-Whitney U test, $p < 0.05$, Benjamini-Hochberg corrected). An asterisk indicates that there is a significant difference in scores between the two sequencing depths.

| | **Bracken** | | **Centrifuge** | | **DUDes** | |
|---|---|---|---|---|---|---|
| Method | Abundance accuracy corrected p-value | F1-score corrected p-value | Abundance accuracy corrected p-value | F1-score corrected p-value | Abundance accuracy corrected p-value | F1-score corrected p-value |
| Medoid | 1.000000 | 1.000000 | 1.000000 | 1.000000 | 1.000000 | 1.000000 |
| GGRaSP | 1.000000 | 1.000000 | 1.000000 | 1.000000 | 1.000000 | 1.000000 |
| MC-0.95 | 1.000000 | 1.000000 | 1.000000 | 1.000000 | 1.000000 | 1.000000 |
| MC-0.97 | 1.000000 | 1.000000 | 1.000000 | 1.000000 | 1.000000 | 1.000000 |
| MC-0.99 | 1.000000 | 1.000000 | 1.000000 | 1.000000 | 1.000000 | 1.000000 |
| GC-0.95 | 1.000000 | 1.000000 | 1.000000 | 1.000000 | 1.000000 | 1.000000 |
| GC-0.97 | 1.000000 | 1.000000 | 1.000000 | 1.000000 | 1.000000 | 1.000000 |
| GC-0.99 | 1.000000 | 1.000000 | 1.000000 | 1.000000 | 1.000000 | 1.000000 |
| SL-0.95 | 1.000000 | 1.000000 | 1.000000 | 1.000000 | 1.000000 | 1.000000 |
| SL-0.97 | 1.000000 | 1.000000 | 1.000000 | 1.000000 | 1.000000 | 1.000000 |
| SL-0.99 | 1.000000 | 1.000000 | 1.000000 | 1.000000 | 1.000000 | 1.000000 |
| CL-0.95 | 1.000000 | 1.000000 | 1.000000 | 1.000000 | 1.000000 | 1.000000 |
| CL-0.97 | 1.000000 | 1.000000 | 1.000000 | 1.000000 | 1.000000 | 1.000000 |
| CL-0.99 | 1.000000 | 1.000000 | 1.000000 | 1.000000 | 1.000000 | 1.000000 |

Table S9: Assessment of statistical difference in outcome per accuracy metric (abundance accuracy, F1-score), per profiler, for bacterial genus-based species-level experiment between 450 thousand readpairs and 5 million readpairs samples (two-sided Mann-Whitney U test, $p < 0.05$, Benjamini-Hochberg corrected). An asterisk indicates that there is a significant difference in scores between the two sequencing depths.

| | **Bracken** | | **Centrifuge** | | **DUDes** | |
|---|---|---|---|---|---|---|
| Method | Abundance accuracy corrected p-value | F1-score corrected p-value | Abundance accuracy corrected p-value | F1-score corrected p-value | Abundance accuracy corrected p-value | F1-score corrected p-value |
| Medoid | 1.000000 | 1.000000 | 1.000000 | 1.000000 | 1.000000 | 1.000000 |
| GGRaSP | 1.000000 | 1.000000 | 1.000000 | 1.000000 | 1.000000 | 1.000000 |
| MC-0.95 | 1.000000 | 1.000000 | 1.000000 | 1.000000 | 1.000000 | 1.000000 |
| MC-0.97 | 1.000000 | 1.000000 | 1.000000 | 1.000000 | 1.000000 | 1.000000 |
| MC-0.99 | 1.000000 | 1.000000 | 1.000000 | 1.000000 | 1.000000 | 1.000000 |
| GC-0.95 | 1.000000 | 1.000000 | 1.000000 | 1.000000 | 1.000000 | 1.000000 |
| GC-0.97 | 1.000000 | 1.000000 | 1.000000 | 1.000000 | 1.000000 | 1.000000 |
| GC-0.99 | 1.000000 | 1.000000 | 1.000000 | 1.000000 | 1.000000 | 1.000000 |
| SL-0.95 | 1.000000 | 1.000000 | 1.000000 | 1.000000 | 1.000000 | 1.000000 |
| SL-0.97 | 1.000000 | 1.000000 | 1.000000 | 1.000000 | 1.000000 | 1.000000 |
| SL-0.99 | 1.000000 | 1.000000 | 1.000000 | 1.000000 | 1.000000 | 1.000000 |
| CL-0.95 | 1.000000 | 1.000000 | 1.000000 | 1.000000 | 1.000000 | 1.000000 |
| CL-0.97 | 1.000000 | 1.000000 | 1.000000 | 1.000000 | 1.000000 | 1.000000 |
| CL-0.99 | 1.000000 | 1.000000 | 1.000000 | 1.000000 | 1.000000 | 1.000000 |

Table S10: Assessment of statistical difference in outcome per accuracy metric (abundance accuracy, F1-score), per profiler, for bacterial strain-level experiment between 450 thousand readpairs and 5 million readpairs samples (two-sided Mann-Whitney U test, $p < 0.05$, Benjamini-Hochberg corrected). An asterisk indicates that there is a significant difference in scores between the two sequencing depths.

| | **Bracken** | | **Centrifuge** | |
|---|---|---|---|---|
| Method | Abundance accuracy corrected p-value | F1-score corrected p-value | Abundance accuracy corrected p-value | F1-score corrected p-value |
| Medoid | 0.18260 | 0.010058* | 0.57075 | 0.84988 |
| GGRaSP | 0.41673 | 0.013315* | 0.57075 | 0.67985 |
| MC-0.95 | 0.96985 | 0.010058* | 0.57075 | 0.80773 |
| MC-0.99 | 0.09158 | 0.024590* | 0.57075 | 0.67985 |
| GC-0.95 | 0.18260 | 0.021794* | 0.57075 | 0.67985 |
| GC-0.99 | 0.09158 | 0.021794* | 0.57075 | 0.67985 |
| GC-0.999 | 0.09158 | 0.021794* | 0.57075 | 0.67985 |
| SL-0.95 | 0.18260 | 0.010058* | 0.57075 | 0.84988 |
| SL-0.99 | 0.18260 | 0.010058* | 0.57075 | 0.67985 |
| SL-0.999 | 0.18260 | 0.010058* | 0.57075 | 0.67985 |
| CL-0.95 | 0.18260 | 0.010058* | 0.57075 | 0.84988 |
| CL-0.99 | 0.32268 | 0.010058* | 0.57075 | 0.67985 |
| CL-0.999 | 0.17518 | 0.024753* | 0.57075 | 0.67985 |

### 8 Accuracy versus reference set size

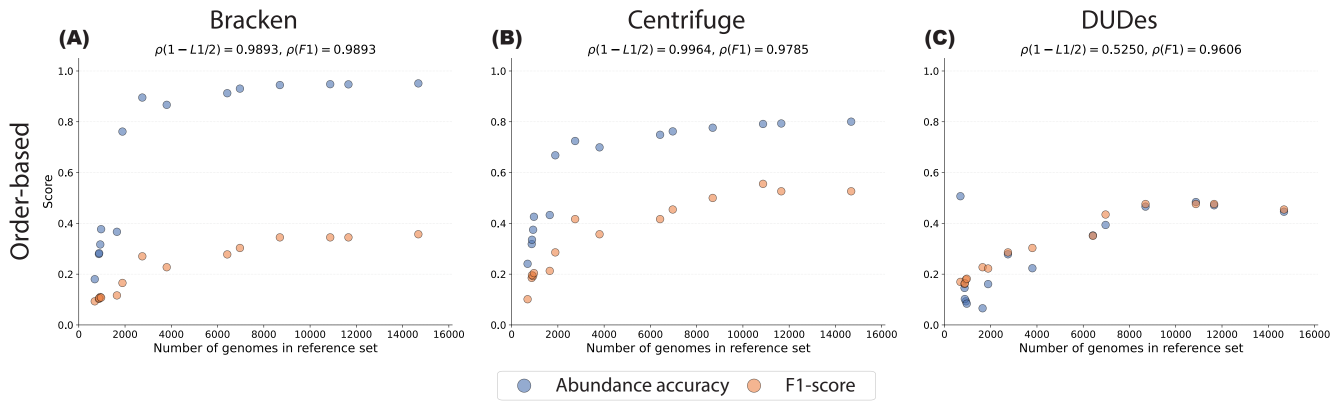

Figure S17: Median accuracy (abundance accuracy, F1-score) of taxonomic profiling tools across bacterial order-based, species-level experiments against the number of genomes in a reference set. Panels A-C are arranged in three columns (profiling tools: Bracken, Centrifuge, DUDes). $\rho$-values in subtitles show Spearman correlaction between number of genomes included and median accuracy score.

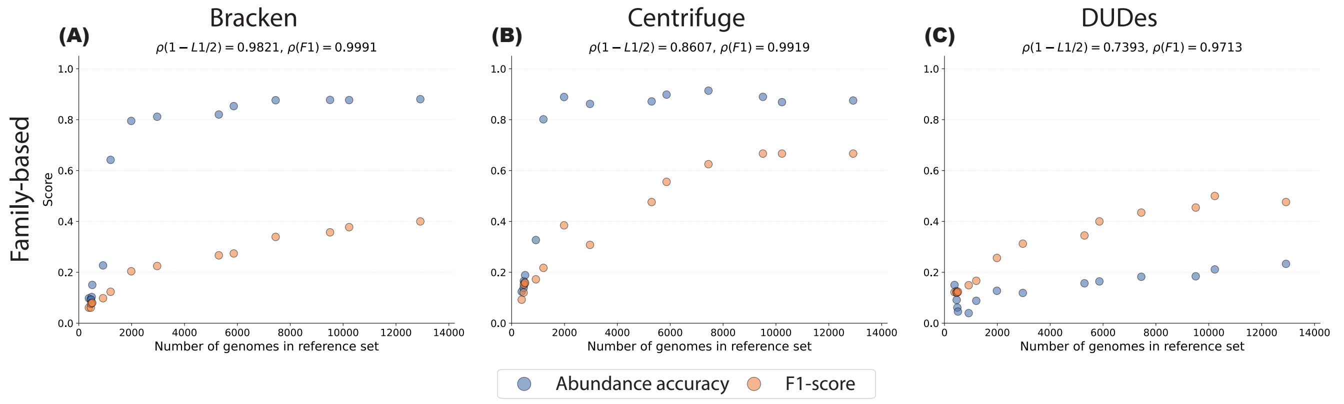

Figure S18: Median accuracy (abundance accuracy, F1-score) of taxonomic profiling tools across bacterial family-based, species-level experiments against the number of genomes in a reference set. Panels A-C are arranged in three columns (profiling tools: Bracken, Centrifuge, DUDes). $\rho$-values in subtitles show Spearman correlaction between number of genomes included and median accuracy score.

### 9 Comparison between real and simulated mock community

To assess the difference in accuracy between the real and simulated mock community samples, we effectively performed a Monte Carlo-like way of deriving p-values for every profiler separately. More specifically, letting $x_1, \ldots, x_{10}$ denote the metric values on the simulated samples for one profiler/method/metric combination, and $x_{\text{real}}$ the corresponding outcome on the real sample, we perform a two-sided test as follows: first, we compute the median accuracy of the simulated values. Denoting the median as $c$, we then quantify the deviations in simulated samples as $d_i = |x_i - c|$, and the deviation in the real sample from the simulated distribution as $d_{\text{real}} := |x_{\text{real}} - c|$. The empirical two-sided p-value was then calculated as:

$$p = \frac{1 + \sum_{i=1}^{10} \mathbb{I}(d_i \geq d_{\text{real}})}{10 + 1},$$

where $\mathbb{I}$ is the indicator function. Afterwards, per profiler and metric, p-values were corrected using the Benjamini-Hochberg procedure, and p-values below $\alpha = 0.05$ were considered significant. Table S11 shows the adjusted p-values for all methods and profilers.

Table S11: BH-adjusted p-values comparing taxonomic profiling accuracy on the real mock community sample against analogue simulated samples.

| Method | Bracken abundance accuracy | Bracken F1 | Centrifuge abundancy accuracy | Centrifuge F1 |
|---|---|---|---|---|
| All | 0.1061 | 0.0909 | 0.0909 | 0.1591 |
| Medoid | 0.1061 | 0.0909 | 0.0909 | 1.0000 |
| GGRaSP | 0.1061 | 0.0909 | 0.0909 | 1.0000 |
| MC-0.95 | 0.1061 | 0.0909 | 0.0909 | 0.1591 |
| MC-0.99 | 0.2937 | 0.0909 | 0.0909 | 0.1591 |
| GC-0.95 | 0.5455 | 0.0909 | 0.0909 | 0.1591 |
| GC-0.99 | 0.1061 | 0.0909 | 0.0909 | 0.1591 |
| GC-0.999 | 0.1061 | 0.0909 | 0.0909 | 0.1591 |
| SL-0.95 | 0.1061 | 0.0909 | 0.0909 | 1.0000 |
| SL-0.99 | 0.1061 | 0.0909 | 0.0909 | 1.0000 |
| SL-0.999 | 0.1061 | 0.0909 | 0.0909 | 0.1591 |
| CL-0.95 | 0.1061 | 0.0909 | 0.0909 | 1.0000 |
| CL-0.99 | 0.1061 | 0.0909 | 0.0909 | 1.0000 |
| CL-0.999 | 0.1061 | 0.0909 | 0.0909 | 0.1591 |
